# Challenges in Undergraduate Synthetic Biology Training: Insights from a Canadian iGEM Student Perspective

**DOI:** 10.1101/2020.11.04.365999

**Authors:** Patrick Diep, Austin Boucinha, Bi-ru Amy Yeung, Brayden Kell, Xingyu Chen, Daniel Tsyplenkov, Danielle Serra, Andres Escobar, Ansley Gnanapragasam, Christian A. Emond, Victoria A. Sajtovich, Radhakrishnan Mahadevan, Dawn M. Kilkenny, Garfield Gini-Newman, Mads Kaern, Brian Ingalls

## Abstract

The last two decades have seen vigorous activity in synthetic biology research and ever-increasing applications of synthetic biology technologies. However, pedagogical research on synthetic biology is scarce, especially when compared to some scientific and engineering disciplines. Within Canada, there are only three universities that formally teach synthetic biology programs; two of which are at the undergraduate level. Many Canadian undergraduate students are instead introduced to synthetic biology through participation in the annual International Genetically Engineered Machine (iGEM) competition where they work in design teams to conceive of and execute a synthetic biology project that they present at an international jamboree. We surveyed the Canadian landscape of synthetic biology education through the experience of students from the Canadian iGEM teams of 2019. Using a thematic codebook analysis, we gathered insights to generate recommendations that could empower future iGEM team operations and inform educators about best practices in teaching undergraduate synthetic biology.

## INTRODUCTION

The development of the synthetic biology field over the last twenty years shares parallels with that of computer science in the 1950s and the internet in the 1990s. The invention of computers and computing technologies marked a turning point in history: electronic machines began to permeate our daily lives, leading to wide-spread social impact and accelerated innovation in numerous disciplines.^1^ Rapid advancements in computer science drove a need for nimble pedagogical computing curricula that was responsive to the field’s extraordinary pace of growth – information could become outdated within the timespan of an undergraduate program. Over multiple phases of development, computing education and its pedagogical foundations adopted inquiry-based learning, collaborative learning, praxis, and situated critical pedagogy.^2–5^ Similarly, engineering education has also evolved in this trajectory over the past half-century.^6–8^

Contemporary synthetic biology (henceforth SynBio) from the early 2000s onward is also heralded as a similarly impactful technology because it is altering how human civilization interacts with the living and its interface with the organic and inorganic worlds.^9–12^ While the definition of SynBio is strongly contended,^13,14^ we provide the following working definition for this report:

> Synthetic Biology is a transdisciplinary field pertaining to the bottom-up creation and top-down augmentation of biomolecular and cellular entities that can be designed from characterized parts and systems. Implementation sciences are integrated into the design and commercialization of these entities to achieve social acceptance through agreed-upon standards and values, to develop policy and biosecurity mechanisms to manage risks, and to safely promote responsible research and entrepreneurial innovation.

A defining feature of SynBio is its collaborative nature where several knowledge domains intersect in biological design and implementation. The extent of these collaborations can be categorized as multidisciplinary, interdisciplinary, or transdisciplinary.^15–17^ While these terms are often used interchangeably, they are not synonymous. Multidisciplinarity refers to several teams working separately towards the same goals. SynBio is predominantly interdisciplinary because of the need for laboratory biologists, computational scientists, and engineers to closely collaborate on the *same* team rather than in separate teams.^18,19^ However, the increasing reliance on ethicists, legal specialists, and social scientists in the SynBio innovation process indicates a trend towards broader transdisciplinarity. Here, individuals in the team begin to possess blends of traditionally disparate knowledge domains (*e.g.* STEM practitioners with non-STEM training such as biochemists with entrepreneurial experience) that allow them to participate in various roles on a team. The integration of these disciplines helps to ensure that considerations needed for social acceptable and commercial feasibility are accounted for earlier in the research pipeline, which radically alters the SynBio solution space.^20–22^ This transdisciplinary nature of Synbio poses a grand challenge for educators in the field: what are the best teaching practices for collaborative synthetic biology education? Many SynBio practitioners and educators have entered the field with specialized disciplinary training. Agapakis (2014) highlights a consequence of this:

> “For conversations between synthetic biology and other design fields, misunderstandings can arise not only over specialist language but also over shared terms that have different meanings in different contexts,” and quoting Alexandra Daisy Ginsberg from the BioBricks Foundation SB6.0 conference, “in art and design, I use the ‘experiment’ as an open-ended process to open up and reveal potential ideas; in science, the ‘experiment’ is a tool to generate data to test a hypothesis.” ^19^

Disciplinary communities have created unique jargon and interpretations of terminologies to describe their theories and concepts, so SynBio educators with only specialized disciplinary training frequently teach SynBio content with a narrower disciplinary lens familiar to them. Without intervention, it can thus be difficult for students to leave such classrooms with an appreciation and respect for other disciplines and their contributions to SynBio problems, as well as an understanding of how to collaborate with others’ whose expertise and knowledge are vastly different, yet relevant. The ability to communicate using the jargon and language of several disciplines is thus essential to present and future SynBio practice that should be taught in undergraduate SynBio curricula. However, the pedagogical literature that outlines such learning objectives is limited, and standards for what constitutes essential SynBio practitioner skills and qualities remain to be defined. As noted by Kuldell (2007),

> “The newness of synthetic biology makes ‘typical’ instruction nearly impossible. For example, how can a teacher properly assess ‘mastery of subject matter’ when the foundational framework and professional competencies of the field have yet to be determined? Effective communication skills and sophisticated reasoning may distinguish experts from novices and so might be considered appropriate readouts for accomplishment, but the measures for success in these areas are imprecise and difficult to apply.” ^23^

Historically, computer science and engineering educators investigated and adopted an array of teaching methods to best prepare students for their future careers. SynBio educators will need to follow a similar strategy to identify what the current pedagogical challenges are, how these challenges change over time due to advances in the field, and draw upon existing pedagogical literature to inform educational experiments that could contribute to developing best practices for teaching SynBio. The current body of SynBio pedagogical literature is small, but has begun to address these needs in piecemeal fashion and provides a basis for further research. SynBio educators have developed teaching tools^24–27^ and initiated dialogue about SynBio pedagogy by discussing the importance of some learning objectives, whereas others have described the implementation and performance of SynBio curricula.^23,28,29^ To contribute to this growing body of literature, this report focuses on gathering insights relevant to undergraduate synthetic biology education in Canada by narrowing in on students participating in the international Genetically Engineered Machine (iGEM) competition, through which the majority of Canadian SynBio education currently takes place.

iGEM is a yearly international SynBio design competition for primarily undergraduate students. To compete in the competition, student teams need to conceive of a SynBio research project, execute it, and meet specific project deliverables. Most deliverables are tied to specific judging criteria that are evaluated by panels of iGEM judges for teams to earn Bronze, Silver, or Gold Medal, which are measures of the quality of the project rather than Special Prizes that can be awarded by judges depending on unique criteria. These deliverables and criteria are often essential to the ultimate goal of iGEM projects, “which is to solve problems pertinent to the world and its inhabitants with the help of SynBio” (igem.org/About). Not only do student teams need to execute on research questions that are biologically and mathematically driven, but they need to address social and ethical questions related to their work by engaging with stakeholders and reflecting on the impact of their project deliverables.

We assessed the learning experiences of iGEM design teams from 17 Canadian institutions using a survey comprised of an electronic (e-)questionnaire and online video call interviews. The goal was to implement a grounded theory approach to generate insights that could be used to improve iGEM design team operations, and to highlight pedagogical approaches that students respond well to for SynBio educators to consider in their planning. In this report, we synthesized five Meta-Themes by integrating our e-questionnaire data with the thematic codebook analysis of the interview transcripts (Table 1). Finally, we concluded with recommendations for iGEM design teams and SynBio educators. Throughout this report, we introduced terminology from psychology and pedagogy to better frame the Meta-Themes and student experiences within published literature with the aim of highlighting future avenues of research in SynBio pedagogy.

**Table 1.** Summary of all Meta-Themes and cognate themes from codebook analysis of interview transcripts

| Meta-Theme | Themes |  | Frequency | Quotes |
| --- | --- | --- | --- | --- |
| 1: Conflict Challenges Social Cohesion | A | Strong social cohesion is an integral part of the iGEM experience | 87%, 13/15 | 1, 4, 5 |
|  | B | Interpersonal conflict was notable and impacted the team dynamics | 67%, 10/15 | 2, 6, 7, 8 |
|  | C | A notable lack of social presence | 20%, 3/15 | 3 |
|  | D | Interpersonal conflict was noted to be a non-issue | 33%, 5/15 | 9 |
|  | E | A code of conduct was implemented within the team's policies | 33%, 5/15 | 10, 11 |
| 2: The Role of Support Network and Host Institution Engagement | F | Teams experienced high levels of support network engagement | 60%, 9/15 | 12, 13 |
|  | G | Teams experienced low levels of support network engagement | 40%, 6/15 |  |
|  | H | Unbalanced engagement with the support network that negatively impacted teams | 33%, 5/15 | 14, 15, 16, 17 |
|  | I | Teams believed their institution perceives them with a positive outlook | 67%, 10/15 | 18, 19, 20 |
|  | J | Institutional engagement was surface-level and needs to be deepened | 40%, 6/15 | 21, 22 |
|  | K | Teams believed their institution perceives them with a negative outlook | 33%, 5/10 | 23, 24, 25, 26 |
| 3: Skills Development and Intra-team Knowledge Transfer | L | iGEM provides opportunities to develop career-relevant skillsets | 100%, 15/15 | 27, 28, 29 |
|  | M | Dependence on intra-team knowledge transfer for training | 93%, 14/15 | 30, 31, 32, 33 |
|  | N | Programs do not prepare students for the iGEM experience | 60%, 9/15 | 34, 35, 36, 37 |
| 4: Training Difficulties with Dry-Lab and Human Practices Sub-teams | O | Absence of formal DL and/or HP sub-team training | 80%, 12/15 | 38, 39, 40, 41, 42, 43, 44, 45 |
| 5: Design Principles for Teaching Undergraduate Synthetic Biology | P | Incoherent synthetic biology curricula | 60%, 9/15 | 46, 47, 48, 49, 50 |
|  | Q | A notable lack of synthetic biology courses offered | 46%, 7/15 | 51, 52, 53 |

## METHODS

This study was conducted using a protocol approved by the University of Toronto Research Ethics Board (#38550). The survey is comprised of (i) an electronic (e-)questionnaire for which consent was electronically obtained, and (ii) recorded online video call interviews for which consent was obtained using scanned physical forms and signatures. All teams and individuals have been anonymized and all data are aggregated; quotes were decoupled from the interviewees to uphold confidentiality.

### Participant Recruitment

In September 2019, we invited 19 Canadian iGEM teams to participate by directly e-mailing student leaders or indirectly contacting them through their teams’ faculty supervisors (*i.e.*, principal investigators and teachers). 2 teams did not respond, so 17 teams proceeded through our survey. Through purposive sampling, we exclusively recruited student leaders to participate in the survey to better describe the overall team experiences. The survey was completed by 2 high school students representing their high school teams, 12 undergraduate students (*i.e.*, presidents, lead coordinators or team representatives), and 3 graduate student advisors that had worked closely with their undergraduate team members.

### Data Collection

The e-questionnaire (Supplementary Materials—Appendix A) was created in October 2019 using Google Forms and distributed via e-mail. The purpose of the e-questionnaire was to gather team-level demographic information that could describe the operating parameters for teams. Specifically, these parameters were sectioned into (i) general background, (ii) support network, and (iii) training resources. The general background section probed basic information about the size of teams and their organizational structure, competition performance, and level of funding. The support network section explored the size of the team’s support network and the extent of their interactions with the team. A support network is defined as the group of faculty supervisors (*i.e.* professors, teachers, administrative staff); instructors (*i.e.* post-doctoral fellows, graduates students) that provide dedicated training, mentorship, and administrative support; and advisors who are similar to instructors, but have less committed roles and usually only assist with troubleshooting issues. The training resource section explored how students were being prepared for the iGEM/SynBio project environment. The e-questionnaire was completed in January 2020.

In December 2019, the interviews (Supplementary Materials—Appendix B) were scheduled via e-mail exchanges with our interviewers and conducted over the subsequent three months using Zoom. These interviews were recorded using Zoom’s built-in recording function. The purpose of the interview was to provide participants the opportunity to expand on some of their answers in the e-questionnaire, and for us to ask more open-ended questions. Congruent with the sectioning of the e-questionnaire, the interview protocol consisted of questions related to general background, support network, training resources. The general background section gathered information about the timeline for projects and the social dynamics of the team. The support network section probed the relationship between teams and their faculty supervisors, as well as for the impressions the iGEM team had on their local student body and faculty body at their host institution. The training resources section probed for information about the student learning experience when working on an iGEM project, and the relationship between this experience and existing academic courses related to SynBio. The interviews were completed in March 2020. Two teams declined interviews. One team responded to our interview questions in written format.

### Data Analysis

The e-questionnaire quantitative data was collected into a spreadsheet through Google Forms and visually analyzed using Microsoft Excel, then graphically prepared for publication using Python 3.6.

The recorded interviews were automatically converted by Zoom to audio MP3 files following completion of the interview protocol. The audio files were manually transcribed to text for thematic analysis with codebooks generated using elements of standard codebook development.^30,31^ Question 1 (Q1) and Q2 (Supplementary Materials—Appendix B) for the general background section were not included in the thematic codebook analysis and were separately discussed (Figure S5). The type of information gathered by Q1 and Q2 was related to the activities that teams were performing throughout the duration of the project, so it was more appropriate to visually analyze the data in a manner that retained the continuity of the different timelines. After generating codebooks for Q3-Q9, we compared their observed themes and found the Cohen’s interrater score (a statistic used to determine agreement in observed themes in thematic codebook analyses) to be at an acceptable level (κ = 85%) for themes that all of our analysts observed with frequencies ≥ 20%.^32^ Observed themes were considered to be reflecting the same phenomenon if they could be clearly combined to synthesize an emergent theme, which we call a “Meta-Theme”, that provides multiple perspectives for a singular phenomenon.

Representative quotes that are used for further analysis have been lightly edited for clarity (using clarification brackets) and brevity (using bracketed ellipses to exclude some text). To protect the privacy and anonymity of participants and their teams, this report does not use team names or pseudonyms. Furthermore, the gender of persons referenced by the interviewees was masked by substituting gendered pronouns with the gender-neutral pronouns, “they” and “them”.

## RESULTS AND DISCUSSION

We depicted the landscape of Canadian synthetic biology education in the following sections by exploring five Meta-Themes based on quantitative data from our e-questionnaire and qualitative data from our thematic codebook analysis of the interview transcripts. We describe these Meta-Themes in rich detail as a basis for further quantitative research, rather than determining the prevalence of specific phenomena and their correlations with each other. Table 1 summarizes each Meta-Themes’ unique set of themes and the frequency which the interviewees commented on each of them. Representative quotes for each theme were used to inductively synthesize the cognate Meta-Theme, which allowed us to connect existing literature with the interviewee’s experiences to draw useful insights for iGEM design teams and SynBio educators. Additional context for basic team demographics, funding information, and a canonical timeline for Canadian iGEM projects are described in Figures S1, S2, and S5, respectively.

### Meta-Theme 1: Conflict Challenges Social Cohesion

We observed the theme “Strong social cohesion is an integral part of the iGEM experience” (87%, 13/15; – Quotes 1, 4, 5), but also observed that “Interpersonal conflict was notable and impacted the team dynamics” (67%, 10/15; Quotes 2, 6, 7). This dissonance as illustrated by Quotes 1 and 2 led us to explore the relationship between social cohesion, defined by Dimas *et* al. (2019) as the integration of the members through emotional bonds and a sense of belonging; and conflict, operationalized by Lee *et al.* (2019) as task conflict (*i.e.,* issues with distribution of resources, procedures and policies, and implementation) or relational conflict (*i.e.,* interpersonal and communication issues).^33,34^

> “Yeah, we tried to create a very social environment. […] A lot of us were already friends from prior courses, and the new people, we just helped them integrate. And often you would just be hanging out with these people because you’d be in meetings with them all the time or in-lab. And then outside, we’d have different get-togethers, dinners…I don’t know if we had ‘parties’, but we definitely went to watch movies and stuff like that.” – Quote 1

> “In the end, […] we didn’t want to take action and sanction the bad behaviors, which kind of hurt us in the development of our team.” – Quote 2

To our knowledge, the interplay between social cohesion and conflict within iGEM teams has not been studied, but abstracting an iGEM design team as a *general* design team enables the use of existing literature to understand this interplay. For example, iGEM design teams are akin to those formed in industry to drive new product development and engineering capstone design teams that are formed for the completion of undergraduate degree requirements. There are many metrics to evaluate the successfulness of these design teams, but organizational psychologists have operationalized the construct of “success” as 1) team creativity and team innovation^35,36^ and 2) team learning ^34,37^, which are both indirectly assessed by the iGEM judging criteria and are therefore linked to competition outcomes.

Team creativity is conceptualized as the generation of novel and useful ideas and solutions; team innovation describes how such advances are implemented through inventions.^35,36^ Rodríguez-Sánchez *et al*. (2017) showed that team creativity is driven by the combination of social cohesion and task cohesion, the latter defined as the shared commitment of members towards common goals.^36^ Vestal & Mesmer-Magnus (2020) found team innovation was shown to be driven by the presence of members with expertise in a variety of disciplines whose knowledge overlap isn’t strongly redundant that can work together well.^35^ These studies make evident the importance of having a strong social environment within a design team for more success in terms of higher creativity and innovativeness.

Team learning can be conceptualized as the process of habitualizing behaviors and performing actions conducive to critical thinking and knowledge application such as seeking feedback, exploring and testing new ideas, and reflecting on misjudgments.^34^ Dimas *et al*. (2019, 2020) applied a nonlinear dynamical system (NDS) approach to model the relationship between team learning, social cohesion, and team culture (*i.e.,* the set of norms, values, and actions that members use to shape their decision-making). Through a cusp catastrophe model, they demonstrate teams can switch between high and low states of team learning based on the social cohesion of teams with the same team culture. Practically, we interpret this as the need for leaders of design teams to carefully modulate the social cohesion of teams to ensure a high state for team learning is maintained – too high and phenomena like groupthink^38^ can occur that inhibits critical thinking; too low and social loafing can occur, which reduces productivity. One theme “A notable lack of social presence” (20%, 3/15; Quote 3) illustrates this issue. Quote 3 shows how poor social membership can also weaken communication channels that can be dually used to strengthen social bonds and discuss project matters.

> “[Human practice members] don’t come often [to social functions] so it was kind of hard to contact them, but we still got through [to them using] emails and stuff. It could be improved.” – Quote 3

While no teams reported excessively high social cohesion, some teams noted the importance of social cohesion for task cohesion (*i.e.,* forces exerted on members to work towards shared team goals) and for lasting social bonds beyond the iGEM competition.

> “If you’re friends with your co-workers, you are more likely to get work done in a much more effective and better manner and that was very much true [from our results].” – Quote 4

> “The people I joined [human practices] with, I would consider those people my best friends at this point in time, and the mentors […] still spoke to them regularly. And so I think that really facilitated a positive experience.” – Quote 5

While the development of strong social bonds was reported in nearly every Canadian iGEM team, many of the same teams reported task and relational conflict. Hurst & Mostafapour (2018), in their study of engineering undergraduate capstone design teams, described eight categories of conflict that they observed.^39^ Particularly relevant to iGEM design teams are the categories poor project management (Quote 6), lack of clarity on team members’ roles and expectations (Quote 6 & 7) and lack of participation and team membership (Quote 7 & 8).

> “One of the [more] responsible people left us as well. […] So, a month before the competition, we had to do it from zero. It was really tiring, […] we didn’t have a solution because it was just put before our eyes, and the only thing we could do was to do the job. And some signs could have lead us to see it before, but we defended this person […] as it was an old member and we were used to seeing a lot of work from this person […] it was hard to accept that it wasn’t the case this year.” – Quote 6

> “Once the summer started we had a breakdown in our team and the wet lab lead decided that [they] didn’t want [their role] anymore and I was heavily counting on [them] to be the primary person on the floor instructing students and I had seen [their] enthusiasm and initiative and [they were] always there. That person you know you can count on, so that for our team was devasting. We really had to pick ourselves up and shake it off and say ’with you or without you we will do this’. And that’s exactly what we did.” – Quote 7

> “[…] someone didn’t do anything, so we wanted to maybe tell this person that ‘you cannot be part of this if you just don’t do anything’, but this person was intimately linked to two others, so it’s really hard when you don’t want to have more authority and you want everything to go smoothly […]. We ended up having the teachers say ‘well, if you don’t answer this by tomorrow, you’re out of the team’ and it pretty much went smoothly. But we still didn’t act fast enough, so it kind of slowed us and we took too much time.” – Quote 8

How conflict influences design team success is a heavily-studied contentious area. Some researchers have reported task conflict to be negatively associated with team creativity^40^, but this has been recently challenged by the argument that task conflict is positively and directly associated with team creativity, yet negatively and indirectly associated with team creativity through relationship conflict that occurs as a consequence of the task conflict.^33^ This research highlights the importance of conflict management to ensure design teams are successful. A theme we observed that sheds light on potential conflict management strategies comes from iGEM teams where “Interpersonal conflict was noted to be a non-issue” (33%, 5/15; Quote 9). One strategy to minimize relational conflict is to prevent it all together.

> “I think this year we went entirely conflict-free […]. We picked people that would work well together. I think that’s a big part and having those social aspects. We did trivia together, fun things outside of iGEM on a weekly basis.” – Quote 9

Vestal & Mesmer-Magnus (2020) argued that higher team relational resources (*i.e.,* knowledge that is acquired by previous teamwork, in which members become aware of the content and applicability of other’s expertise) can increase team innovation by compensating for the knowledge gaps that individual members may have.^35^ In other words, teams with high relational resources have worked together several times before, so they understand how to use their strengths to cover their teammates weaknesses. However, iGEM design teams regularly undergo turnover due to the nature of the competition being an annual event, so student leaders may only be able to recruit some members from the previous generation’s team that have worked together. One approach to counter this loss in relational resources is noted in a study by Takai & Esterman (2017) where they encouraged instructors during group activities to “assemble team members who have similar preferences on work structure” and contended “team members collectively have to have diverse expertise in order to complete the required tasks but at the same time team members have to be similar enough to work collaboratively.” ^41^ This research suggests that design teams can be more strategic with their recruitment by evaluating their applicants on individual criteria *and* considerations for the likelihood of them socially integrating into the team community based on personalities and work-style preferences. Individual technical expertise and research experiences are necessary for task cohesion, but the teams’ breadth of expertise and anticipated social cohesion could be addressed before the project starts.

To handle conflict when it does arise, we observed the theme that “A code of conduct was implemented within the team’s policies” (33%, 5/15; Quotes 10, 11). Codes of conduct (CoCs) are documents used to provide clear guidance in situations where the best line of action is not clear, as illustrated in Quotes 2, 6, and 8. CoCs explicitly state the values of the team, outline roles and responsibilities, delineate what ethical professional conduct should look like, and alert members to the consequences of certain actions and poor behaviors to encourage a healthier work environment. Quote 10 highlights the importance of CoC agreements being a consensual process that is intended to be used as a “best practices” document that which iGEM students can refer to.

> “So when everyone joins the team, at our first team-wide meeting, we have a code of conduct that we go over with everyone. They all get emailed it and they have to sign and return it. […] it kind of just goes [over] our expectations: how you should be behaving, things that are allowed, not allowed, […] what you do if you feel like you need to bring something up with someone.” – Quote 10

> “We did draft a constitution for our team, a new thing this year, to try to build in conflict resolution into our processes and this included a three-strike approach where we all agreed that this is the system we would be using. […] After the three strikes, a member would have a bad standing designation and we would have to consult with our supervisors, our PIs, which would let them know that the privileges of being in an iGEM team like, for example, a fully-funded participation in the Jamboree, or even attending the Jamboree, were contingent upon them fixing their issues and making up for any contributions that they missed, but they were not kicked off the team.” – Quote 11

Quote 11 exemplifies how CoCs can be used as an instrument to provide objectivity when subjectivity obscures the best line of action. While conflictive situations are nuanced and should ideally be dealt with on a case-by-case basis, CoCs still provide student leaders and their supervisors insurance – a signed, agreed-upon, multi-staged protocol to deal with varying degrees of misconduct and grievances. Developing CoCs can be difficult because foreseeing future types of conflict that may arise in the existing and future generations of the iGEM design team is hard. Slimani *et al.* (2007) provide a useful taxonomy of conflicts to learn about what can arise. Supplementary Materials—Appendix C provides for a CoC template that iGEM design teams could adapt to their needs based on this taxonomy.^42^

Taken together, the research drawn to explore the interplay between social cohesion and conflict seen in iGEM design teams strongly suggests that the former is needed for various success metrics – creativity, innovation, learning – and the latter does present a challenge to design teams, but can be prevented through better recruitment decisions and mitigated through CoCs that categorize conflict and offer protocols to address them.

### Meta-Theme 2: The Role of Support Network and Host Institution Engagement

iGEM design teams have support networks comprised of three types of individuals: faculty supervisors, instructors, and advisors (*cf*. Methods—Data Collection for definitions). The institutions that host these iGEM design teams also provide several forms of support. Our thematic codebook analysis yielded a range of themes associated with these external parties supporting the teams. Here, we first discussed the role of support network individuals after reflecting on e-questionnaire data along with experiences of iGEM design team students, then in similar fashion discussed the role of host institutions. Finally, we consolidated these discussions by building a conceptual model to better understand the relationship between the support network, host institution, and the iGEM design team.

#### Support Network: Faculty Supervisors, Instructors, and Advisors

Our e-questionnaire evaluated the accessibility, helpfulness, and involvement of the support network individuals from the perspective of iGEM design team students (Figure S3). Instructors and advisors scored higher than faculty supervisors for accessibility and helpfulness, possibly due to their more frequent meetings with the teams and stronger involvement in the development of the project. However, faculty supervisors received mixed scores. They were mostly somewhat accessible and somewhat helpful or less, which could be due to their less frequent meetings with the teams, and contrasts with instructors and advisors. We also found that most faculty supervisors had limited or no involvement with project development and again the opposite to be true for instructors and advisors. These apparently negative observations for faculty supervisors before commencing interviews indicated to us that there might be an important relationship between the frequency of their interactions with teams, their involvement in the development of the project, and success metrics that deserved further analysis.

We observed “teams experienced high levels of support network engagement” (60%, 9/15; Quotes 12, 13), which contrasts with the opposite theme where “teams experienced a low levels of support network engagement” (40%, 6/15).

> “[The faculty supervisors] weren’t very hands-off, but they let the students and the grad students drive the project […] they were still there for funding authorization, the signing of safety forms, and administrative duties as they were essential for that […] when we were super stuck with what we wanted to do or nothing was working, we would consult with the [the professors] during the weekly meetings” – Quote 12

> “So we have one professor […] who has been either the main PI for the team every single year, and if [they’re] not the PI, [they’re] still involved. And the way [they’re] mainly involved is either providing some of [their] own lab space to us, and attending weekly meetings to give feedback on what we’re doing and to help with troubleshooting. [They] give advice on what we should do next and also [provide] us with a project code at the university, so when we apply for internal funding […], the grant funds go directly into our research account.” – Quote 13

More teams reporting notably higher engagement with their support network in the interviews corroborated our e-questionnaire data (Figure S3) that is left-skewed towards higher scores for accessibility, helpfulness, and satisfaction with meeting frequencies. However, the level of support network engagement is not dichotomous as evidenced by the theme for “unbalanced engagement with the support network that negatively impacted teams” (33%, 5/15; Quotes 14-17) that shows it is a spectrum for levels of engagement. Quotes 14 and 15 illustrate excessive levels of engagement from the student perspective, and Quotes 16 and 17 starkly illustrate the opposite.

> “We did have conflicts with our graduate student members. That was an issue from [before] and the reason why we wanted that formal [conflict resolution] process […]. They’re advisors because they train us and give us counsel on how to actually conduct the project. But it ends up being the case that they’re always involved and always there every meeting, getting involved in fundraising, administrative things like safety forms and communicating with [the] PI. […] Because of such involvement with the progress of the project, personal issues get involved and that’s why we tried to implement this more formal structure” – Quote 14

> “There were a few issues with the supervisor and certain people. [The] supervisors at one point lost the goal of iGEM which was that it was a student-run project, and they started making all the calls. And so, a few people on the team got a little irritated by that. And the supervisor actually made the call about who would go to Boston without consulting anybody and a lot of people were a little annoyed by that too because the students were not consulted. And it’s something that we can’t handle super well because it’s a supervisor […].”– Quote 15

> “So our [professor] had an idea of what [they] wanted to do and we basically split the parts together. So for this year, it wasn’t really a collaborative project. […] it was hard to start because we basically tried to do different parts of a project. […] doing an assembly is one thing, but my part was basically verifying if it’s possible to clone […] different inserts into the backbone and trying to incorporate that into yeast, so it’s such a big jump. It doesn’t connect well. I would say it’s like separate projects. And I was verifying if it [works] in yeast, and someone else was doing Gibson, and another person was verifying Type II-S. So, I think it was more scattered and that was a problem because we couldn’t really communicate between each other.” – Quote 16

> “There was a time when our [professors] were very, very busy and they weren’t coming to as many [weekly] meetings and I felt as though it was definitely a disadvantage to our team when our [professors] stopped coming to the meetings because as much as the graduate students have this independence, we don’t have as much knowledge for troubleshooting as they do.” – Quote 17

It is clear that faculty supervisors, instructors, and advisors all play important external roles for the iGEM design teams based on the e-questionnaire data we collected, but it appears from the interview data that faculty supervisors have uniquely profound effects on the success of teams that cannot be simply reduced to a demand for their expertise and administrative powers. For example, Quote 13 illustrates how the faculty supervisors provide teams with stability; Quote 15 speaks to the need for faculty supervisors to promote autonomy; Quote 17 speaks to the role faculty supervisors have in building students’ competence. The faculty supervisor while being backstage for most of their role has an impact that is spotlighted by their team’s performance. We therefore hypothesize faculty supervisors play a special role in nurturing strong team learning environments that can enable further creativity and innovation – ultimately influencing the success of teams.

#### Host Institutions

The e-questionnaire acquired team funding information that could help gauge the level of engagement between host institutions and iGEM design teams (Figure S2). Most funding came from the lab or faculty supervisor that was supporting the team and additional funding from different levels within an institution further categorized as departmental, faculty, and university-wide sources. It is unclear whether these funding sources are stable or not over a 3-year period due to most scores spread between “somewhat unstable” and “somewhat stable”. It is clearer that teams are generally dissatisfied with the amount of funding they have, which we speculate is due to the increasing costs of team registration ($4000 CAD) and Jamboree registration ($1100 CAD per attendee) in the 2019 competition year. However, 14/17 teams received at least $10,000 CAD in funding, which would allow teams to register for the competition and send a few students to the Jamboree in Boston. While there could be more funding, institutions are clearly supporting iGEM teams from a financial perspective; the interview data corroborates this notion. Regarding how teams think their host institution perceives them, we found the theme that “teams believed their institution perceives them with a positive outlook” (67%, 10/15; Quotes 18-20), and suggests this funding is tied to the institution recognizing the value of students participating in the iGEM competition. This is especially clear in Quote 20 where the interviewee outlines the value propositions of students participating in the iGEM competition.

> “I think the university as a whole has a favourable look on the iGEM team, more so with the Faculty of Science. We’ve been featured by their faculty many times over, and in their media releases, interviews, and in fundraising campaigns for the faculty, we’ve been featured with our project and some of our members. The Faculty of Science has also contributed the most financially to the team. […] They haven’t given us support in wet lab space, so we had to rely on our PIs to get that lab space for us.” – Quote 18

> “We’ve gotten a lot of good feedback from the university. […] We’ve gotten letters of support and congratulations from the Dean of Maths and Sciences, from the Dean of Research, and even from the librarians at our university, so in that regard, iGEM is definitely looked at in a good stance. […] And even the [newspaper would] do little tidbits of stories for us […] so I think it’s all positive.” – Quote 19

> “I think at our school we are lucky to have a really good reputation. I think we’ve always done well at the competitions, so it looks good for the university as well. Plus, the university uses us as advertisement because we are an opportunity for undergraduates to get experience in the wet lab throughout their degrees, so it looks really good on the university to have that, or to offer that experience. […] Plus, its interdisciplinary, so it’s not just that we’re offering this experience to […] a small number of people; almost anybody at [our institution] can apply to be a part of the team. […] it’s just easier for us to get internal funding because people are aware of the program.” – Quote 20

However, funding is only one form of support that some host institutions can provide iGEM design teams. As Quotes 18 and 19 illustrate, media attention and validation from institutional leadership are other forms of support that are important for connecting these teams to the grander community at these institutions. The theme “institutional engagement was surface-level and needs to be deepened” (40%, 6/15; Quotes 21, 22) suggests teams need multiple forms of support from their host institutions.

> “[…] we’ve gotten a lot of support in terms of ‘oh my god that’s a great project, so cool that students are doing this’. […] one of my goals personally as I joined the leadership team over the past few year and sort of trying to push our team to the next level, has been trying to translate this into additional funding.” – Quote 21

> “I would just like to have a little bit more support in general from the university. A lot of professors in our university, we can’t go ask for help because they don’t like the [iGEM] program. So, it’s not like we can just go and ask different faculty members for help if we need it because they don’t fully support the program.” – Quote 22

Quote 22 begins by acknowledging the host institution does support the team in some capacity (*i.e.* funding), but later expresses the desire for faculty members at the institution to appreciate the iGEM program more and be more willing to support the team. We suspect this sentiment is common across several iGEM design teams because it is corroborated by the theme that “teams believed their institution perceives them with a negative outlook” (33%, 5/10; Quotes 23-25).

> “I do remember a few conversations that people would be a bit hesitant about iGEM because they wouldn’t think that it would actually come up with a tangible solution; it would more so be a science project that wouldn’t go anywhere. That’s really what we wanted to convince them against, because we wanted to say that this is actually something that can address real life problem that we can solve with synthetic biology and this innovative approach that we are pitching.” – Quote 23

> “[…] every now and then there are a few professors that are like ’that’s not real research, it’s not the same as if you were in a lab doing primary research’, but they’re very few and far between.” – Quote 24

> “You hear mixed reviews, like some people think it’s valuable, [and] some think it’s a waste of time. It’s more of the people that don’t see the value in gold medal requirements more than anything. That seems to be a big debate, like ‘cool you got a gold medal that really means nothing’.” – Quote 25

> “[The advisors] opinion is ’Okay, that’s cool that you’re doing this.’ But they don’t really care that much. But at the same time, I do think they get annoyed sometimes when we ask them so many questions like ’What are we actually doing in the lab?” – Quote 26

We propose this negative outlook of iGEM stems from a misunderstanding of the intended goals of the competition. iGEM projects involve conducting academic research, experiential learning, and stakeholder engagement exercises. They can thus simultaneously act like research groups, educational programs, and start-up incubation space. The goals of the competition for iGEM projects are therefore to have some combination of publishing peer-reviewed papers, learning the skills necessary to execute a successful project, and addressing issues informed by stakeholder engagement. Combining these activities may not be perceived by all as fitting within institutional mandates, yet iGEM projects can produce well-rounded critical thinkers to complement their more specialized training in academic labs, discipline-specific courses, or start-ups they may embark on. The ultimate consequence of these negative outlooks is the alienation of students on the iGEM design teams from the grander research community, as exemplified through Quotes 23-26. We therefore hypothesize that institutional leadership from faculty members plays a critical role in nurturing strong team learning environments that can enable further creativity and innovation – ultimately influencing the success of teams.

#### A Conceptual Model and Tools for Improved Engagement

Previously, we hypothesized that both faculty supervisors and institutional leadership can ultimately influence the success of iGEM design teams by affecting the team’s learning environment, and therefore their creative outputs and innovation. The mechanisms underpinning how this influence happens are not reported in literature for this specific context, so we sought to develop a conceptual model to first understand how faculty supervisors may influence team success, then extended this model into simple engagement and diagnostic tools for teams to systematically address engagement challenges they may be encountering with both faculty supervisors and their host institutions to increase their chances at success.

We began by invoking self-determination theory (SDT) to delineate the relationship between faculty supervisors, host institutions, and iGEM design teams. SDT describes the cognitive mechanisms underpinning human intrinsic motivation (*viz.*, in the absence of external factors like financial compensation or material rewards).^43^ It postulates that a person can perform an activity or adopt a behavior through two lines of reasoning known as introjection or integration; the former involves one doing something whose purpose they do not identify with or accept, and the latter involves one doing something for the opposite reason. SDT proponents argue that integration leads to *self-determination* where “one does the behavior wholly volitionally because of its utility or importance for one’s personal goals.” ^43^ The three psychological needs contingent for self-determination are autonomy, competence, and relatedness. Autonomy refers to the feeling of control over one’s actions and the sense that one’s action are aligned with their own values and identity.^44^ Competence refers to the feeling that one is efficacious in their environment.^45^ Relatedness refers to the feeling that one is connected to and belongs to a community.^45^

We contend a requisite for nurturing strong team learning environments is the fostering of self-determination in iGEM design team students. Faculty supervisors play a role in fostering self-determination because of their potent ability to satisfy students’ psychological needs for autonomy and competence since they hold the most expertise and administrative power compared to instructors and advisors. To satisfy these needs, faculty supervisors need not maximize their engagement with teams, but find an optimal level of engagement. We define this *engagement* as 1) the *frequency* of interactions, which is related to psychological need for competence; and 2) the *influence* they exercise in changing the direction of the iGEM project, which is related to the psychological need for autonomy. Frequency is related to competence in this conceptual model because, according to our interview data, interactions between teams and faculty supervisors typically involve troubleshooting, mentorship, coaching, and similar activities that improve the knowledge and skills that students have that help them be efficacious in their project environments. Influence is related to autonomy because the more opportunities students have to make important decisions about the direction of the project, the more control students may feel over their project environment. Frequency of interactions and influence in decision-making thus form the two axes of the conceptual model that we call the Design-iGEM Team Oversight (DiTO) diagram (Figure 1). While this discussion continues with a focus on faculty supervisors, the DiTO diagram and further discussion are generalizable to all individuals in the support network (*i.e.*, instructors and advisors too).

**Figure 1.**
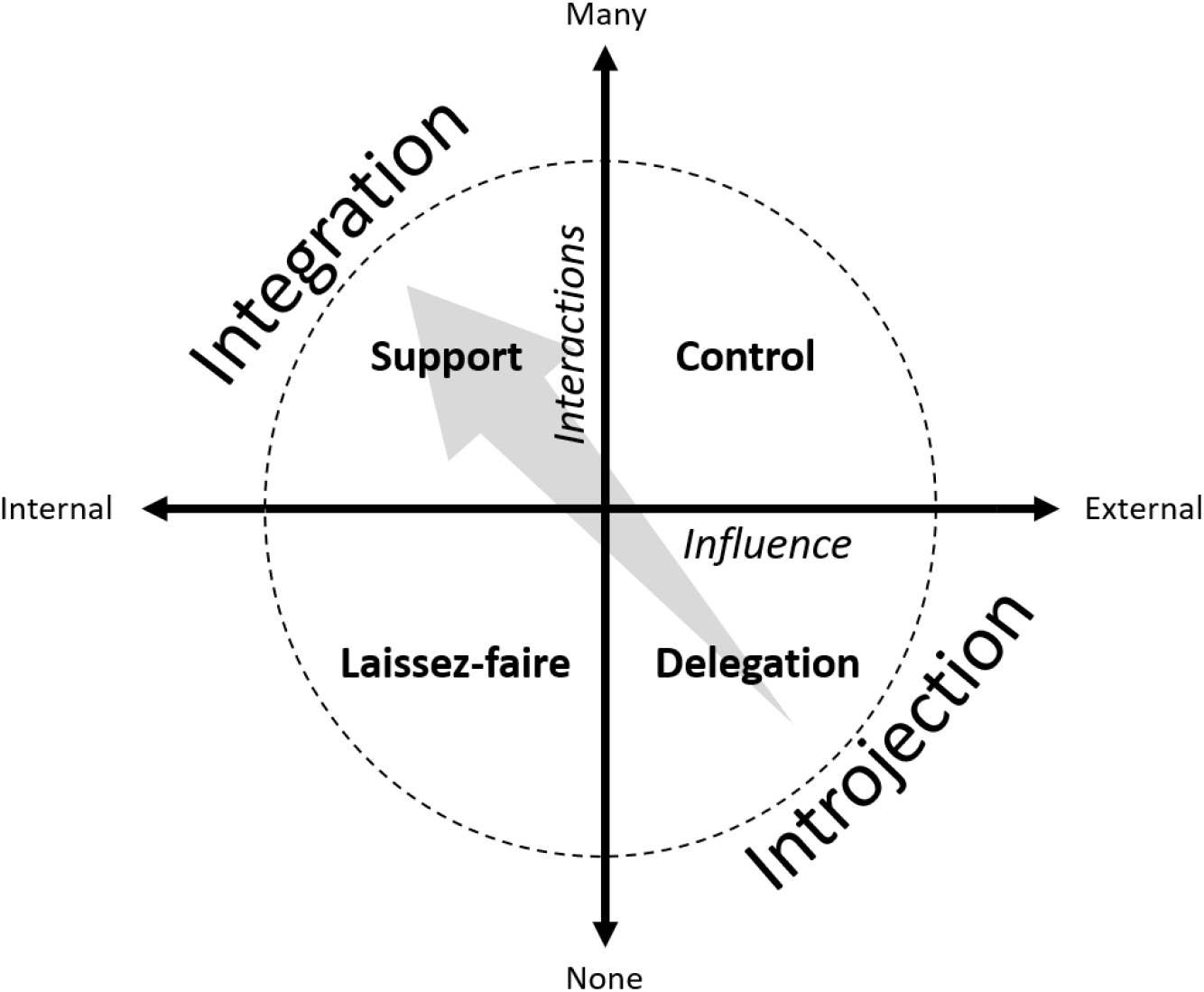
Design-iGEM Team Oversight (DiTO) Diagram. Engagement strategies can be characterized in terms of two parameters: 1) the influence the support team individual has on decisions related to project ideas and progress from internal (with the team) or externally (x-axis) and 2) the frequency of interactions with the team from none to many (y-axis). The quadrant labels are the four types of engagement strategies: Support, Control, Laissez-faire, and Delegation. The arrow signifies a spectrum of introjection to integration across the quadrants.

The DiTO diagram is comprised of four quadrants representing different engagement styles. The top-left quadrant describes the scenario where the faculty supervisor frequently interacts with the team in a manner that empowers the team to hold most of the decision-making power. This style is called *Support*, exemplified by Quotes 12 and 13, and is known to promote team creativity and overall success.^46,47^ The bottom-right quadrant describes the opposing scenario where the faculty supervisor bears most of the decision-making power over the project, yet has minimal interactions with the team. This style is called *Delegation*, exemplified by Quote 16. The top-right quadrant describes a scenario where teams experience many interactions with the faculty supervisor, yet the team holds less decision-making power. This engagement is called *Control*, exemplified by Quotes 14 and 15. Finally. the bottom-left quadrant describes the scenario where teams experience minimal interactions with the faculty supervisor, but the team has more decision-making power. This style is called *Laissez-faire*, exemplified by Quote 17. We view introjection and integration as two ends of a spectrum spanning from Delegation engagement styles to Support styles; this is indicated by the gray diagonal arrow in Figure 1.

We contend Delegation styles are the least conducive for fostering self-determination in students because the faculty supervisor is the least engaged in both frequency of interactions and how often they empower the team to make their own decisions that ultimately define the trajectory of their learning. Although introjection found in Delegation styles might be ubiquitous, we suspect it could exacerbate student mental health in competitive environments like iGEM. Cooper *et al.* (2020) conducted a study that sought to connect literature on depression with the lived experiences of undergraduate student researchers in the United States.^48^ They reported students feeling that they could not explain their personal situations to their research mentors because they “weighed the consequences of their research mentors’ displeasure against the consequences of revealing their depression and decided it was not worth admitting to being depressed.” Further, they reported that students experienced difficulty when faced with experimental failure, and those living with depression were particularly susceptible to triggering of depressive symptoms due to failure. Faculty supervisors are not mental health practitioners, but Cooper *et al*.’s study connects the iGEM-derived DiTO diagram to the more general topic of low faculty supervisor engagement with their undergraduate researchers and its connection to poorer student performance.

Potentially compounding the problem of Delegation engagement styles is the disconnectedness some interviewees expressed between their team and host institution (Quotes 22, 23, 35). While most host institutions provided sufficient funding to allow iGEM design teams to register for the competition and send students to the Jamboree, it is evident that there is also a demand for host institutions to be more involved with the teams and to take a proactive role in helping student leaders nurture a stronger team learning environment. This demand is related to final psychological need in SDT to complete this conceptual model: relatedness. Institutional leaders, such as deans and departmental chairs, have the potent ability to increase the sense of belonging in iGEM design teams because they can most easily champion the value propositions for having students participate in the iGEM competition and rally support given their leadership role among professors and research staff.

Combined, faculty supervisors and host institutions are hypothesized to influence the success of teams by playing a role in nurturing a strong team learning environment. This deserves further research. The DiTO diagram is a conceptual model that can be used as starting point for teams and faculty supervisors to discuss engagement styles. Additionally, we offer Supplementary Materials—Appendix E as a roadmap with engagement tools to promote autonomy and competence, as well as Supplementary Materials—Appendix F as a similar road map to promote relatedness. Finally, for teams to better understand the state they are in regarding the quality of their support network and their relationship with their host institutions, we designed a simple scoring rubric (Supplementary Materials—Appendix D) for teams to better understand their circumstances.

### Meta-Theme 3: Skills Development and Intra-team Knowledge Transfer

Participation in the iGEM competition is both co-curricular and extra-curricular because the skillsets acquired by students can complement their programs’ curricula and be applied to real-world work scenarios that are not necessarily the learning goals for programs. We report a consensus theme that “iGEM provides opportunities to develop career-relevant skillsets” (100%, 15/15; Quotes 27-29).

> “Competing in iGEM is a stressful process, but learning how to best talk to others especially when you are stressed, how to stay together as a team and view yourself as a key part of that team and to work to best make sure that everyone is succeeding together… rather than having one person who is doing really good in this one area and one person who is struggling a bit in another area but to bring everyone together and be at a point where everyone can stay uplifted and to know that if one person is struggling you have someone else that is able to help you. There is a lot about team dynamic that you learn along the way.” – Quote 27

> “Being able to ask for help [and] not feeling uncomfortable communicating with mentors or other grad students. Many students spoke of how before they used to be introverted in their studies and after iGEM they are speaking to many throughout faculty and student bodies, and feel much more confident.” – Quote 28

> “And then there is also leadership, not only from the leads themselves but also from the members, because we want to be able to give students the opportunity to do independent research; they need to be able to build up the confidence to be a leader without actually [being] given the title because you can be a leader without being called a leader. Being able to take that role, to take initiative, those are the kind of skills that we really try to allow students to improve upon and give the chance for students to improve upon.” – Quote 29

However, this theme contrasts with e-questionnaire results (Figure S4). We assessed which extra-curricular skills teams had formal training for and discovered most teams did not have formal training for every skill we included in the e-questionnaire like leadership, interpersonal communication, and project management. This suggests iGEM design team students were mostly acquiring these skills through experiences working on the project. More surprisingly, we found most teams indicated they would be interested in formal training for these skills, which highlights an unmet demand that can be understood through the lens of Bourdieu’s concept of social capital.^49,50^

Kovalchuk *et al*. (2017) explored how co-curricular activities were sites where students could slowly accrue and transform cultural-educational capital (*i.e.,* knowledge, skills, and experiences) and social capital (*i.e.,* connections and relationships) into economic capital (*i.e.,* improved job prospects, employment). Since iGEM is both co-curricular and extra-curricular, students have several chances to generate economic capital because they can apply what they learn from class in multi-faceted iGEM project activities destined to be presented to an international audience where their networks can be expanded. Some argue that over-committing to co-curricular and extra-curricular activities can negatively impact students’ GPA scores as it could distract them from their academic work, but the reverse has also been argued: these activities can bolster students’ understanding of course material by reinforcing it through other types of learning in Bloom’s learning taxonomies.^51,52^

In a related near-consensus theme, we observed there was a “dependence on intra-team knowledge transfer for training” (93%, 14/15; Quotes 30-33), which illustrates the co-curricular nature of iGEM design teams since students can learn material they would normally learn in academic courses through direct coaching by senior students in the team, instructors, and advisors.

> “We do have training in the very beginning, but that was with people from last year who did some lab work and they just went through stuff. Like ’this is what this is…’ […] and just learn the basic skills.” – Quote 30

> “[the students] go through the different protocols that have been designed by the upper year students, go over how to do things, how to develop a project, how to develop protocols, how to look at past protocols and sort of adapt it to what they need to do, and building timelines.” – Quote 31

> “Okay, so the way we train them [in] two steps really. The first step is reading and meetings to bring them up to speed on the theory of what we’re doing. So, we typically send them articles and reviews and we discuss them in detail at our meetings and once we feel that they have a good background of the theory of what we’re doing, then we bring them into the lab, and then we do experiments. […] And then once we see that they’re more confident and comfortable with those more simple tasks, then we bring them up to more difficult experiments such as cloning using primer design, Gibson assembly, etc. The way we kind of structure it is always training in theory and via the computer and then once they have a good grasp on that, we let them shadow us in-lab and then after they shadow us a couple times, we then allow them to work independently and then we just are there for any questions or guidance.” – Quote 32

> “[…] we made an entire curriculum; we made slides and we lectured twice a week. We also had a biweekly lab that had lab reports due for that as well, and there was a midterm, there was a final. It was a course for sure. […] we would cover things like iGEM project design, iGEM requirements, but we would also cover like heavy molecular biology topics. […] there was also a dry lab component to that course, but […] they didn’t know what part of dry lab would fit into the project, so they did basic general training on Arduino and coding.” – Quote 33

Quote 33 is unique in that the team developed a well-planned SynBio curriculum for their students with a concerted effort to develop instructional material and assessment tools that were meaningful for their iGEM project. We hypothesize teams that created varying degrees of SynBio curricula needed to do this because of the observed theme that “programs do not prepare students for the iGEM experience” (60%, 9/15; Quotes 34-37).

> “So it really depends on where they’re coming from and their backgrounds. Generally speaking, if we step back from technicalities, in terms of the process that iGEM requires, in terms of project development, execution, collaboration with other teams, preparing for an international competition, do students come in well-prepared? No, I don’t think so. Like, unless they’ve done something similar before, it’s pretty new, and so some of that process takes time to adjust to.” – Quote 34

> “To be quite frank […], when they do the undergraduate lab components, I feel as though we coddle them a lot, so often times when we put these upper-year undergraduate students on the benchtop, they’re not very confident in their abilities. And so, they’re not always prepared to be independent, even at the level of third- and fourth-years.” – Quote 35

> “The professor in charge was working on the pathways of diseases; so we did biosensors and we designed primers. It was synthetic biology in the methods, but [fundamental] in the aim. […] in the end, the research itself is kind of fundamental, but not in the same domain as [synthetic biology].” – Quote 36

> “I wish there was more [synthetic biology] integrated into our current biology classes because right now they are trying to change the curriculum, but at least when I was taking it, it was still very much basic and repetitive between our courses. […] it would help for them to take out more ’old biotech’ so they can update it basically to what is currently being done.” – Quote 37

Quote 35 comments on how undergraduate laboratory courses may not sufficiently train students through inquiry-based learning. Students from these courses may not be well-trained enough in handling failure, troubleshooting experiments, and critically interpreting data so they are therefore not prepared for research activities like those on iGEM teams. Quotes 36 and 37 highlight similar issues (*cf.* Meta-Theme 5—Design Principle 3) where synthetic biology is taught strictly in the context of the professors’ research topic that is often not explicitly synthetic biology, or it is taught as a biotechnology course with a focus on the developments of the 1970s to 1990s. Consequently, the students are not exposed to the richness of contemporary synthetic biology which iGEM teams are embedded in.

In all, these quotes in Meta-Theme 3 illustrated how iGEM projects facilitate the training of co-curricular and extra-curricular skills that better prepare students for their future career paths, which often occurs “in-house” within the iGEM design team. However, this training could be more formalized to ensure students uphold higher standards for such skills. Faculty supervisors play a role in nurturing strong team learning environments that can specifically facilitate the training of these skills (*cf.* Meta-Theme 2), but these individuals are professors and research staff with competing priorities. Cartile *et al.* (2019) observed that “co-curricular activities are not recognized by their host universities for academic credit” in most Canadian settings, and “faculty and staff receive no recognition or incentive toward supervisory involvement.” ^53^ If it is within the institutions mandate to produce more leaders, shifting the development of these skillsets from the periphery (*i.e.*, programs like the iGEM competition) to program curricula (*e.g.*, course-integrated leadership development) may yield better results.^54^ Where this is not possible, increasing incentives for faculty and staff to engage more with iGEM teams could serve both the institution’s and students’ interests.^53^

### Meta-Theme 4: Training Difficulties with Dry-Lab & Human Practices Sub-teams

Based on the e-questionnaire data, most teams divided their work across sub-teams focused on wet-lab (WL), dry-lab (DL), and human practices (HP) topics (Figure S1). The interview data indicated most of the training within the iGEM design team was for the WL sub-team and less frequently for DL; only two teams had formal training for HP. This dominant theme, “absence of formal DL and/or HP sub-team training” (80%, 12/15; Quotes 38 – 45), suggests that teams might be facing difficulties with finding and recruiting students with programs related to DL and HP topics. They may also be having difficulties with training their students in DL topics like modelling biological systems and computational simulations; or HP topics like stakeholder engagement, policy analysis, and economic feasibility studies due to a lack a relevant expertise on the team or amongst the support network.

#### Dry-Lab Training

> “For modelling, our team is probably the weakest in [it]. […] So, since we had no expertise in that at all, we had to enlist the help of […] other Canadian iGEM teams that we collaborated with to help us get trained for that, or just get some more information and understanding of modelling for biological systems.” – Quote 38

> “One of our advisors [has] a lot more experience with modeling than anyone else on the team, so [they] helped a lot with that. More recently, it is definitely an area that we struggle a lot with because we don’t have the resources at our university to be able to do an extensive mathematical model.” – Quote 39

> “It was kind of sad for the sole [dry-lab person] because [they] were reading a lot of books, just basic science. [They] could have gotten more mentorship […] because we don’t have anyone else who knows how to code. They had prior knowledge of how to code, but it’s still difficult” – Quote 40

> “The math team has always struggled to train their members because a lot of them will have such different backgrounds; some of them will be really strong in math, but have never taken a biology course in their life, or [vice versa]. And then often our math leads are not the most organized people in the world, and struggle to give structure to their in training.” – Quote 41

Instruction in mathematical biology, which is fundamental to DL activities in iGEM teams, faces challenges rooted in institutional barriers and disciplinary divides.^55–57^ While there are exceptions, Chiel *et al*. (2010) explained, “biology students and faculty have a different way of ‘knowing’ than students and faculty in mathematics, physics, and engineering.” ^56^ At the level of most undergraduate education, rigorous proofs are the basis of math-focused disciplines while empiricism is the basis of life science disciplines. We hypothesize the disciplinary background of team leaders and their mastery of related skills will influence the ability of the team to train its students in WL, DL, and HP topics. Extending this notion, if a team is strong in its WL capabilities because there are disproportionately more team members and leaders that have a disciplinary background in WL-related fields, then there will be issues with training DL and HP topics and performing project work that requires related skills.

This is true for some teams (Quotes 38-40), but organizational issues may also be a contributing factor to difficulties in training students in any topic (Quote 41). We suspect this training issue posed by this hypothesis might be worsened by Delegation or Laissez-faire engagement styles (*cf*. Meta-Theme 2) due their lower meeting frequencies with the team, but more research is needed to substantiate this claim. If the team leaders intervene by specifically recruiting strong members and leaders in DL-/HP-related fields, or seeks help from other teams with complementary strengths, this issue may be ameliorated (Quotes 38 and 42).

> “[…] there is no official training for [dry-lab and human practices]. What we do with all of our team leads – we start the project [and] we make a Gantt chart. So we have […] on a daily basis a plan of what exactly we want to do, what we want to accomplish. And I think that really sets the precedent because the outreach […] really changes depending on your project, and so does the dry lab stuff, especially for the dry lab, that’s something the directors usually don’t know a lot about, so I’d say they are the most independent unit of the team. Because we cannot help them do a mathematical model or […], I guess for our project, the design for the biosensor. Like, we know nothing about designing a prototype, so it was mostly on them to come up with their own weekly schedule, and we just went through it together to make sure we know what they’re doing” – Quote 42

#### Human Practices Training

> “For the HP team, I don’t think they have any sort of formal training. […] they kind of work on projects and everyone knows what’s kind of going on, but a lot of them don’t have a good understanding of the actual biology of the project. Sometimes the lab leads or members will have a meeting where both of the sub-teams will get together and members can teach each other about what they are doing […].” – Quote 43

> “I wouldn’t call it training because it’s not ‘educational’, but I would call it more ‘informational’. It was [mostly] the people who were more experienced with human practices work [informing] the new recruits [about] what we were doing and what we have done as a team and past iGEM teams, and just giving more context and understanding for human practices but we didn’t have any training sessions, like lab sessions, for human practices.” – Quote 44

The iGEM competition defines HP as the “study of how [the] work affects the world, and how the world affects [the] work” (2020.igem.org/Human_Practices), and serves as the basis for medal criteria that define the scope of iGEM design team projects. While this definition elegantly describes the intent of conducting HP-related work, it does not delineate the knowledge domains needed to effectively conduct such work. Corroborating this notion are Quotes 43 and 44 that represent a common sentiment among interviewees who expressed uncertainty about the expectations of their own HP sub-teams. We contend this general definition of HP can preclude formal training activities and instruction by obfuscating which disciplines need to be drawn on to conduct the HP-related work, and therefore the learning objectives based on these disciplines are undefined. We therefore sought to generate a more precise definition of HP by first examining the literature since the exploration of how science and technology (S&T) impacts society is not new.

Technology assessment (TA) was a concept that existed in the 1970-1990s that involved public actors providing feedback about emerging S&T activities like biotechnology. Ethics, legal, and social implications (ELSI) programs were later popularized in the United States in the 2000s as embedded features in large-scale scientific projects that additionally addressed the ethics and legal matters associated with new S&T. Finally, responsible research and innovation (RRI) was popularized in the European Union in the 2010s and was a strategy to govern S&T developments such that TA and ELSI were embedded earlier in the innovation pipeline to be a driving force that shaped how the S&T was translated to society. RRI contrasts with TA and ELSI because it is proactive rather than reactionary in nature.^58^ From this, we defined the goals of WL and DL-related work to be the development of the S&T, and the goal of HP-related work to be the assessment of the social acceptance and commercial feasibility of the S&T. While mathematical modelling and laboratory experiments are the cornerstones of DL and WL-related work, respectively, communication projects are the cornerstone of HP-related work because RRI requires one to engage with various stakeholders that need different communication skills. One team in our interviews touched upon this succinctly:

> “And then human practices, a lot of it is training in communication, so written and oral communication, especially argumentative communication if that makes sense. So, how to make an argument in a way that speaks to the facts, as opposed to just sort of saying something based on anecdotal evidence and things like that. And then a lot of the training comes into the principles of community engagement and what that looks like, what we can and can’t do, and the importance of thinking about what are approaches to community engagement.” – Quote 45

Given the purpose of HP is to assess the social acceptance and commercial feasibility of S&T related to SynBio, it is clearer which disciplines contribute knowledge and skills needed to accomplish these goals; which include sociology, ethics, policy, risk management, engineering economics, and entrepreneurship. We contend RRI is a useful framework for structuring training activities and educational programs. Betten *et al*. (2018) found the use of application scenarios (*i.e.,* grounded speculations) and techno-moral vignettes (*i.e.,* imagined futures) appeared to be effective at fostering qualities desirable for RRI practitioners in iGEM students: anticipation, inclusivity, reflexivity, and responsiveness.^59^ To our knowledge, the use of RRI to help structure SynBio undergraduate education does not exist in Canada, but deserves further research to ascertain its pedagogical utility. We later continue this discussion of using RRI to understand HP better by combining RRI with business training to re-brand HP as Implementation Sciences (*cf.* Meta-Theme 5—Design Principle 2).

### Meta-Theme 5: Design Principles for Teaching Undergraduate Synthetic Biology (SynBio)

Formalized SynBio curricula in Canada include undergraduate programming at Western University and the University of Ottawa, and the SynBioApps graduate program at Concordia University (Figure 2). Additionally, we found at least 50 undergraduate and graduate courses across Canadian universities that focused on synthetic biology, biotechnology, systems biology, and genetic engineering (Supplementary Materials—Appendix G). In our e-questionnaire, many teams also reported the availability of academic work placements in laboratories with SynBio-related projects, and to a lesser extent industry work placements at SynBio companies. It was clear that SynBio was taught at Canadian institutions with opportunities to apply this knowledge in both academic and industrial environments, but our interview data offered a finer-grain assessment of the quality of the undergraduate SynBio courses. Therefore, we sought to generate design principles for SynBio educators in planning their curricula, instruction, and assessments using the team feedback collected in our interviews.

**Figure 2.**
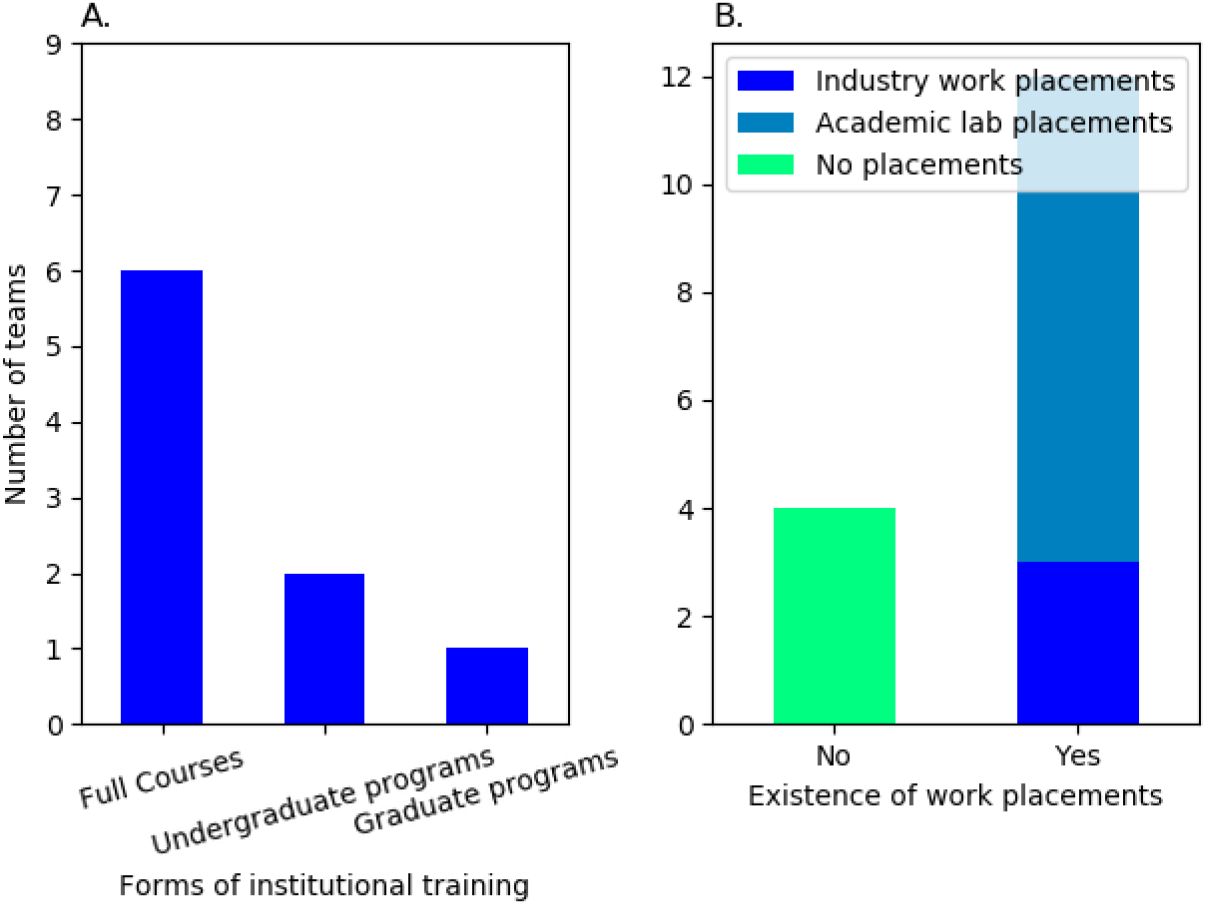
Presence of SynBio learning opportunities at Canadian institutions. Left-panel: availability a formalized academic SynBio courses and programs. Right-panel: availability of SynBio-related academic lab and industrial work placement.

We began by noting common trends in this feedback during the thematic codebook analysis. The first theme was the presence of “incoherent synthetic biology curricula” (60%, 9/15; Quotes 46 – 50) that reported by teams at their host institutions. We defined “incoherent” as the lack of course structure (Quote 46) and issues with the sophistication of the course material (Quotes 47 and 48).

> “I think having something more structured would be more beneficial for us. Ideally, we would have the locally developed synthetic biology [workshop programs] as an actual course in the timetable so students could take it […] for a semester and build up that knowledge, and then be able to be part of the team. […] When it isn’t as structured, we have students who unintentionally miss out on learning one part, which leads to further confusion.” – Quote 46

> “There is […] one biology course called biotechnology in society that talks a little bit about synbio, but almost as a side note in one of the lectures […] and that’s really the extent that people are exposed to synthetic biology.” – Quote 47

> “Maybe also incorporating a third-year synthetic biology course because I feel as though synthetic biology is very particular and although there’s a lot of components that you acquire in other biology courses, I feel that synthetic biology is very precise. So, maybe doing an introductory course to synthetic biology specifically, and then the people that take the third-year course will be better prepared for the fourth-year, more independent course. I don’t know…see, this is the problem…because we get a lot of students wanting to enroll in the fourth-year course, but from what I’ve discussed with my supervisor he says that a lot of the students just don’t fully grasp the articles that they’re reading, so having maybe a third-year introduction to synthetic biology would be beneficial.” – Quote 48

Some teams also reported project-based SynBio courses existed at their host institution that shared parallels with how iGEM projects were conceived of, however these courses were at the fourth-year level for senior undergraduate students that iGEM design teams typically don’t recruit, or the course was inaccessible to most students on the team due to the pre-requisite courses being available only to students in a specific faculty that most of the iGEM design team’s students were not from. This suggests there are sometimes barriers to learning synthetic biology associated with the seniority of the students (*i.e.* too early in their program to access the course), or institutional constraints (*i.e.* lack of cross-faculty SynBio courses).

> “[…] the course kept going up until this last year and it involved students learning about advanced molecular biology techniques and biotechnology, and was just a survey course of different things that are new and emerging in the field of synthetic biology. […] at the end of that, they were responsible for making a project in the same fashion as an iGEM project and then they would present that. […] For 2019 in particular, no one came from that course and no one had taken that course from the time that they were recruited.”– Quote 49

> “[…] they essentially do what iGEM does from start to finish [and] have to come up with a project and the different aspects of the project. So that’s pretty much the extent to what I know about it. There wasn’t any consultation with our team.” – Quote 50

We also collected feedback for how SynBio should be taught from the theme of teams reporting “a notable lack of synthetic biology courses offered” (46%, 7/15; Quotes 51 – 53). Overwhelmingly, students emphasized the need for SynBio courses to teach more than *just* WL-related topics by highlighting the importance of DL and HP-related training (*cf*. Meta-Theme 4). We interpreted this as a demand for more collaborative SynBio courses.

> “I would hope they would include not only the basic fundamentals of synbio in terms of the biology behind it, but also the mathematical components involved in it as well because there is a lot of math in synbio that biology students don’t realize. And engineering components as well. Those basics and as well interdisciplinary type courses. I don’t know how to properly describe that but in other words courses that are not strictly biology, chemistry, or mathematics, or engineering, but mixes engineering mentality into a biology mannerism and vice versa. Even including aspects of humanity and ethics. […] SynBio is very interdisciplinary in the first place as a subject field so you can’t just teach it in a way where it is just one particular subject.” – Quote 51

> “[…] I’m kind of apprehensive of the whole idea of having synbio courses or programs […] because synbio is so multidisciplinary and there are so many things going on, and it can be applied in so many different ways, and I think that if we tried to put it in this box that almost ruins it. I like that I can work with health students and I can work with math students, and we can all bring something and work on a project. […] I definitely think that having SynBio courses would be great, maybe things that can cover techniques in synthetic biology, what’s happening in synthetic biology, or ethics in synthetic biology. […] open to all faculty, not just science or not just engineering.” – Quote 52

> “I think an interdisciplinary course between the Faculties of Engineering and Science and Medicine, that would be very nice. Like, a multi-instructor course where you get different perspectives from the pharma side, from the biotech side, from the industrial microbiology side…I think that would be really interesting where they don’t focus on the particular procedures, but more so…See, I’m an engineering student so I had no idea that we have such granular control on DNA now, and that with even more improvements on the horizon like automation and data science, that we have so much more tools available for bioengineering […] I think making it more accessible for more people is more important and also a project component would be very important for this because it would help the students to engage with each other and that’s one of the coolest things in iGEM is that you get to talk to different disciplines and learn from them, understand what their perspectives are. And human practices, I feel is one of the most important lessons I’ve learned from iGEM are these considerations – considering the needs, wants, and ethical implications to your stakeholders. I think that is the most beneficial lesson that we can gain from these experiences as it’s applicable to every field that you go to. Like, I think it’s super valuable. So, something that has interdisciplinary focus, project work, and value for human practices…I think that would be a really good course for synthetic biology” – Quote 53

Taken together, we extracted four design principles for teaching undergraduate SynBio based on teams’ feedback and further discussed them below: 1) An Interdisciplinary Science Framework and 2) RRI and Business Training (*cf.* Quotes 51 – 53 & Meta-Theme 4), 3) Learning Progression (*cf.* Quotes 46 – 48), and 4) Project-based Situated Learning (*cf.* Quotes 49 & 50). Integration & Implementation Science (I2S) is an emerging discipline that is dedicated to studying the collaboration between disciplinary experts and societal stakeholders tackling complex issues like organized crime, inequitable healthcare, climate change. I2S practitioners are therefore specifically skilled in the integration sciences, the study of how traditionally disciplinary silos can connect their unique approaches to addressing a dimension of a complex challenge; and the implementation sciences, the study of how innovations from different disciplines can be developed with social acceptance and commercial feasibility in consideration.^60,61^ Given SynBio research and iGEM design team projects frequently tackle complex challenges, we propose these design principles can be configured to illustrate their interrelatedness by using the I2S ideology, embodied by the integration sciences and implementation sciences, to scaffold together the four design principles into a potential teaching strategy for undergraduate SynBio (Figure 3).

**Figure 3.**
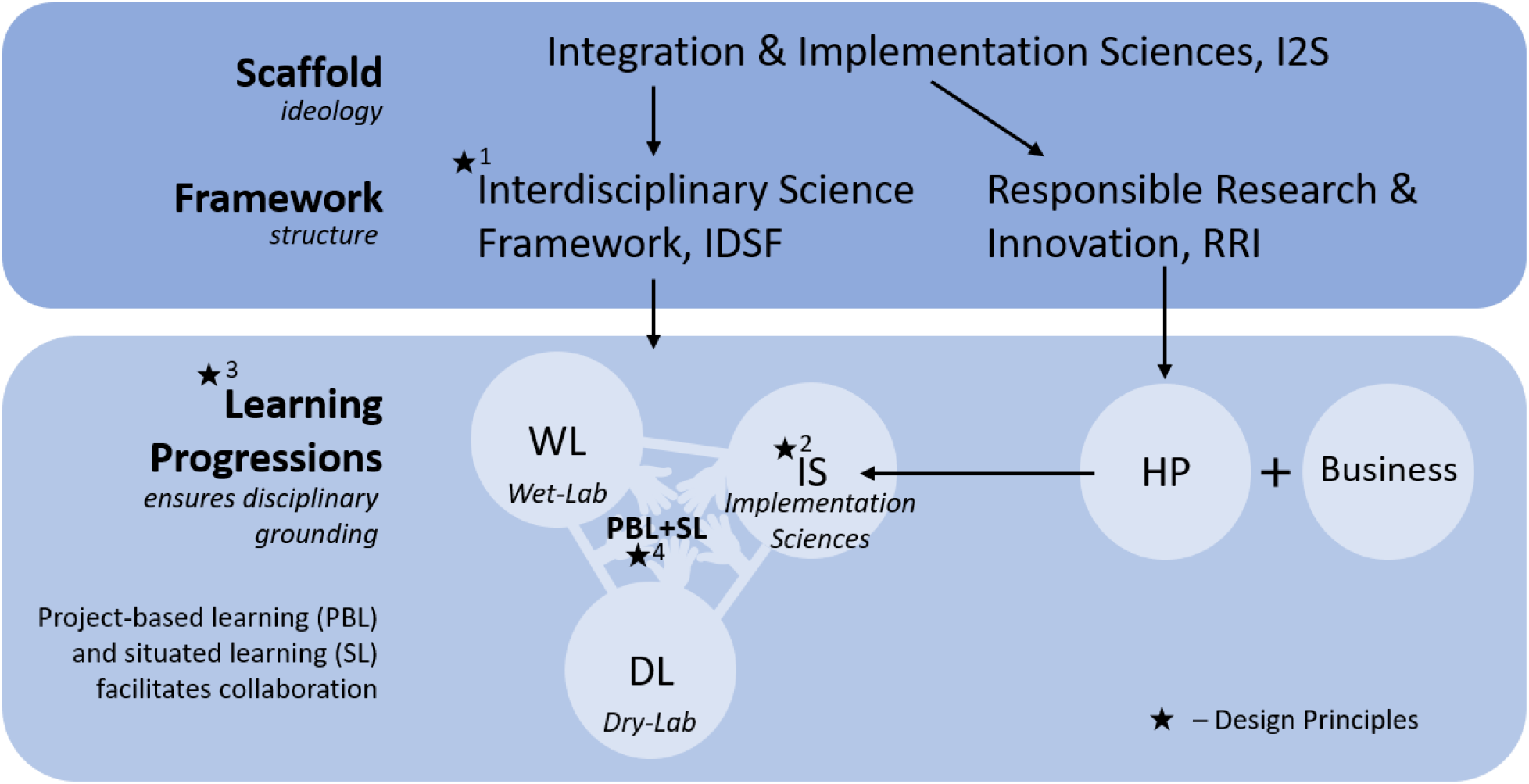
A conceptual configuration of the four major design principles for teaching undergraduate synthetic biology (SynBio). Integration & Implementation Sciences (I2S) provides a pedagogical scaffold for constructing the SynBio learning experience. To implement I2S, the Interdisciplinary Science Framework (IDSF, Design Principle 2) and the Responsible Research & Innovation (RRI) framework are proposed. The combination of RRI with business skills is used to re-brand the traditional work of HP to Implementation Sciences (IS, Design Principle 3). Learning progressions (LPs, Design Principle 1) are used to create long and short-term learning objectives and outcomes for WL, DL, and IS students. To foster an interdisciplinary learning environment, IDSF and project-based situated learning (PBL+SL, Design Principle 4) serve to nurture and improve students’ collaborative qualities and skills.

#### Design Principle 1: An Interdisciplinary Science Framework (IDSF)

de Lorenzo *et al*. (2019) contended SynBio can play a role in addressing some United Nations Sustainable Development Goals (UN SDGs) through bioproduction, remediation, and pollution control.^62^ Addressing these complex environmental and societal issues requires interdisciplinary collaboration between disciplinary experts and societal stakeholders, and therefore draws on integration sciences. To prepare SynBio students for collaborative environments, as is clearly in demand based on our interview data (*cf.* Quotes 51-53), we propose the use of the Interdisciplinary Science Framework (IDSF) described by Tripp & Shortlidge (2019) as the basis for Design Principle 2 because it outlines how students from different programs can be trained to effectively collaborate with each other.^63^ Teaching undergraduate SynBio should therefore implement the four IDSF elements: 1) disciplinary grounding, 2) awareness of different research methods and methodologies, 3) advancement through knowledge integration, and 4) collaborative skills with other disciplines.

Disciplinary grounding refers to the acquisition of *discipline-specific* expertise – a starting point that prevents students from becoming diletantes.^64^ After achieving some mastery of this expertise, students can also begin acquiring *provisional* knowledge of other disciplines: a “working fluency” allowing them to appreciate and utilize different research methods (*i.e.*, tools, equipment, and instruments) and methodologies (epistemological rationale governing disciplines) beyond the traditional scope of their disciplinary expertise. Advancement through integration then requires students to learn to piece together knowledge from disparate disciplines to generate novel ideas and solutions, so students need to be trained to think creatively about how methods or methodologies from other disciplines could be applied to the same challenge they are tackling with their own disciplinary grounding. Finally, students need to be trained in the practice of collaborating with individuals with different disciplinary groundings, which involves learning to frame problems through different disciplinary lenses and understanding others’ roles in the innovation process.

Bridging these elements together is the nurturing of disciplinary humility in students, embodied by inclusive practice, respect for other disciplinary cultures, and reflexivity regarding the limitations of one’s own knowledge, skills, and personal biases.^63^ Tripp & Shortlidge (2019) propose disciplinary humility can be fostered in students by facilitating collaborative group projects addressing simulated real-world problems (*cf*. Meta-Theme 5—Design Principle 4). They have also provided a rubric to illustrate how these IDSF elements could be assessed in a course.^65^

#### Design Principle 2: Responsible Research & Innovation (RRI), and Business Training

As discussed previously (*cf.* Meta-Theme 4—Human Practices Training), the unclear definition of HP used by iGEM design team students was refined through the use of the RRI framework to delineate which disciplinary groundings are needed to contribute to the social acceptance and commercial feasibility of new science and technology (S&T) in SynBio. We propose, as Design Principle 3, the formalization of this re-imagined HP as the second component of I2S: Implementation Sciences (IS). To do this, we suggest explicitly combining RRI training (*i.e.*, sociology, ethics, policy, risk management) with business training (*i.e.*, engineering economics and entrepreneurship). This is analogous to undergraduate engineering programs offering economics courses for engineering students to be formally taught to connect their technical mastery with knowledge and skills that enable the use of this mastery in real-world applications. Design Principle 2 is therefore the development of learning progressions dedicated to preparing SynBio students to assess the social acceptance and commercial feasibility of the S&T they are developing (discussed next in Design Principle 3). These learning progressions could include biotechnology risk assessments, science communication strategies, market research, stakeholder surveying, calculating cost-benefit analyses, and initiating start-ups.

#### Design Principle 3: Learning Progressions (LPs)

A learning progression (LP) is a sequence of stages through which students progress as they build mastery of a concept or skill, and draws on cognitivist and constructivist learning theory. LPs are mostly applied by educators in the planning of learning objectives and outcomes (herein termed learning *goals*) for a series of courses (*i.e.* stages) over semesters and years of a program.^66–68^ They have been popularized in the sciences, and in particular biology, because of their systematic approach to creating courses spanning years of education that have interconnected course learning goals to specifically produce strong scientists and biologists.^69^ All disciplines face difficulties in structuring their teaching strategy and SynBio is not unique, however SynBio pedagogy as an area of research and formalized SynBio curricula at Canadian institutions are still nascent and emerging (*cf.* Figure 2). We therefore argue LPs should be used proactively *now* in the development of new SynBio teaching strategies at Canadian institutions rather than using LPs reactively to student criticism about the unstructured nature of future SynBio courses and programs. Our interview data has already shown this criticism beginning to bud as more undergraduate SynBio courses and programs emerge in Canadian institutions.

A simple grand learning goal for SynBio programs could be to produce strong synthetic biologists. LPs can thus serve as roadmaps for how students’ initial ideas about key SynBio topics (termed a *lower anchor*) mature over multiple courses and throughout the program towards a more sophisticated professional level of practice (termed an *upper anchor*). By defining what this “professional level of practice” looks like based on the expertise of faculty members and potentially feedback from relevant industry stakeholders, SynBio educators can design courses with interconnected learning goals to provide students with structured learning experiences to help them achieve mastery of indispensable SynBio concepts and skills. Scott *et al.* (2019) list five features of science LPs: 1) they consider the breadth of students’ non-scientific colloquial explanations of scientific concepts (lower anchors), 2) they describe student thinking about “big ideas” in science that is related to 3) how students reason when engaged with scientific concepts, 4) they identify what the grand learning goals are to help guide course-level learning goals to systematically train students, and 5) they align assessment tools with broader learning goals of the program with targeted course learning goals to provide reasonable and rigorous benchmarks and milestones throughout the entire program.^69^ To illustrate a SynBio LP, we constructed an LP for genetic circuits (Supplementary Materials—Appendix H).

#### Design Principle 4: Project-based Situated Learning

Design Principle 4 connects Meta-Themes 1, 2, and 3 to the design of undergraduate SynBio programs by complementing theoretical training with project-based learning (PBL) and situated learning (SL). PBL and SL can provide students opportunities to apply their knowledge in collaborative group settings to learn to address complex SynBio problems.

PBL is a form of active learning that draws on project-based learning, and focuses on students combining tackling problems in a long-term team-oriented setting. Educators may involve industry professionals tackling real problems to help frame student projects in the real world, thereby increasing the authenticity of the PBL activities. PBL is related to the Self-Determination Theory (*cf.* Meta-Theme 2—A Conceptual Model and Tools for Improved Engagement) because educators must provide students with some level of project decision-making freedom (autonomy), and balance group work with didactic teaching to ensure students acquire relevant knowledge to effectively tackle the projects (competence). Educators must also create an environment where students feel comfortable asking questions, taking risks, adopting a growth mindset, and be available to give students prompt feedback (relatedness). Proponents of PBL argue the use of projects simulates real-world work experiences more than other forms of teaching like didacticism and behavioral learning, and can therefore function as a career preparation too since students need to learn to manage conflict (*cf.* Meta-Theme 1) and practice other non-technical skills needed for real-world work environments (*cf.* Meta-Theme 3).^70,71^

SL frames the learning process as a socialization process where students acquire knowledge and expertise through “legitimate peripheral participation”, the process of apprenticing someone or some group and learning how to think and behave like those in the same community. This is related to the frequency which students can interact with SynBio educators and synthetic biologists actively researching and working in the field (*cf.* Meta-Theme 2—Support Network: Faculty Supervisors, Instructors, and Advisors). Through this process, one gradually, learns the culture of the community and becomes an active member, which signals that learning has been achieved.^72,73^ For example, multiple iGEM teams reported the use of mentorship to train new members by having them shadow senior members of the team until they became independent enough to perform laboratory work and computations on their own (*cf.* Meta-Theme 3).

## CONCLUSIONS

To our knowledge, this report is the first of its kind to comprehensively survey the majority of iGEM teams within a country to better understand the country’s state of undergraduate SynBio education. Given the exploratory nature of our questions, we probed teams through an e-questionnaire and video call interviews. In the e-questionnaire, we were able to compare teams to each other by collecting general information about their student composition, their feelings about the support network and institutional support, and their access to SynBio education and training. In the interview, we further probed the same team representatives on many topics including their experiences in the competition, team dynamics, training activities, and feelings about SynBio courses and programs that existed at their institutions. A thematic codebook analysis of these interview transcripts revealed over a dozen themes, from which five salient Meta-Themes emerged.

Using a grounded theory approach that drew on literature in psychology and pedagogy, the Meta-Themes were able to generate recommendations to improve iGEM design team operations, and for SynBio educators to consider when designing projects, courses, and entire programs (Table 2). While there are clear limitations to this study (Supplementary Materials—Appendix I), our report lays some of the foundations for the field of SynBio pedagogy. We hope this report sparks further discussion about undergraduate SynBio education and helps to drive new interest in SynBio pedagogy. Our research group plans to continue exploring many of the unexplored research avenues noted throughout this report, beginning with the prototyping and testing of SynBio LPs for WL, DL, and IS topics through a virtual learning environment we are developing for undergraduate students.

**Table 2.** Summary of recommendations to iGEM design teams and synthetic biology educators

| Meta-Theme | iGEM Design Team Recommendations | Synthetic Biology Educator Recommendations |
| --- | --- | --- |
| 1: Conflict Challenges Social Cohesion | future conflict can be mitigated during recruitment by considering the applicants' fit into the social community in addition to their technical merit<br>use of a Code of Conduct (CoC) to navigate difficult situations with unclear resolution paths. Appendix C provides a CoC template for teams to adapt | conflict management strategies should be taught to improve students' abilities to maintain the strong social cohesion that is necessary for collaborative work environments<br>educators should be available to help intervene in situations where an authority figure is needed |
| 2: The Role of Support Network and Host Institution Engagement | teams should be aware the engagement style (*cf.* DiTO diagram) of individuals in their support network and their host institution's engagement can greatly influence their success. Appendix D offers a useful diagnostic tool to help teams evaluate the quality of these relationships<br>Appendix E provides a pathway to help teams improve engagement with their support network, particularly faculty supervisors; Appendix F offers a pathway to help teams strategize how to better engage the institution | promoting self-determination in students is key to having strong team learning environments. This can be done by empowering students with *autonomy* (*i.e.*, giving them more decision-making power), *competence* (*i.e.*, sufficiently coaching them to be efficacious in their project environment), and *relatedness* (*i.e.*, ensuring they are member of a larger supportive community)<br>when supervising teams during group projects, a Support engagement style naturally promotes self-determination |
| 3: Skills Development and Intra-team Knowledge Transfer | seek leadership and project management workshops for undergraduate students facilitated by the host institution or external parties to strengthen team functionality<br>formal mechanisms like transition documents can help ensure the next generation of iGEM students learn from the previous year's mistakes and reflections | a cornerstone of iGEM projects from a training perspective is the improvement in students' leadership, teamwork, communication, and project management skills. The design of SynBio course and programs should involve the training of students to have these qualities that contribute to their preparation for collaborative work environments |
| 4: Training Difficulties with Dry-Lab and Human Practices Sub-teams | teams should consider advertising their recruitment to departments and faculties with students specifically trained in DL and HP-related topics to strengthen these dimensions of their iGEM project<br>rather than rely on HP medal criteria to define the HP sub-team's activities, their activities should revolve around the social acceptance and commercial feasibility of the S&T that the WL and DL sub-teams are developing | *This section includes recommendations based on both Meta-Themes 4 & 5 due to their relationship to SynBio pedagogy.*<br>the design of SynBio workshop series, courses, and programs should incorporate four design principles:<br>1. IDSF: this framework outlines a strategy to promote collaborative skills in students regardless of their home in WL, DL, and IS-related disciplinary groundings<br>2. RRI and Business Training: combined, these constitute a re-imagined dimension of SynBio, Implementation Sciences (IS) focused on assessing and improving the social acceptance and commercial feasibility of new S&T<br>3. LPs: a strategy to structure the learning goals of multiple courses to ensure students reach upper anchors that will prepare them for careers in SynBio<br>4. PBL+SL: active learning strategies that incorporate insights from Meta-Themes 1, 2, and 3 |
| 5: Design Principles for Teaching Undergraduate Synthetic Biology | for more collaborative team environments, student leaders should consider fostering disciplinary humility through increased communication (and socials) between WL, DL, and HP sub-team members<br>HP sub-teams should consider re-branding themselves to IS sub-teams whose (well-defined) work in an iGEM project is to use RRI and business skills to assess and improve the social acceptance and commercial feasibility of the S&T the team is developing |  |

## ACKNOWLEDGEMENTS

We thank Rebecca Allen for assistance with recruiting team representatives for the e-questionnaire and survey, Katariina Jaenes for assistance with preparing the research ethics board application and interviews, James Bagley for assistance with interviews, and Alexandre Tremblay for assistance with identifying synthetic biology courses in Québec. We thank our advisory board for their contributions to the review of this manuscript, especially Brian Ingalls for his extensive feedback in earlier drafts. Finally, we thank the Canadian 2019 iGEM design teams for participating in this research that we anticipate will drive advances in the field of SynBio pedagogy.

## CONTRIBUTOR’S STATEMENT

P.D. – Conceptualization, Methodology, Investigation, Formal Analysis, Writing (Original, Review & Editing), Project Administration; A.B. Conceptualization, Methodology, Investigation, Writing (Review), Project Administration; B.A.Y. – Investigation, Visualization, Project Administration; B.K. – Formal Analysis, Visualization, Writing (Original, Review); X.C. – Formal Analysis, Writing (Review); D.T. – Investigation; D.S. – Investigation; A.E. – Formal Analysis, Visualization; A.G. – Formal Analysis; C.A.E. – Writing (Review); V.A.S. – Writing (Review); R.M. – Project Administration; D.M.K. – Writing (Review); G.G.N. – Writing (Review); M.K. – Writing (Review); B.I. – Writing (Review & Editing)

## REFERENCES

1. Danziger, J. N. Social Science and the Social Impacts of Computer Technology. Soc. Sci. Q. 66, 3 (1985).

2. Zachary, J. L. An introduction to computing for engineers: new approaches to content and pedagogy. in Technology-Based Re-Engineering Engineering Education Proceedings of Frontiers in Education FIE’96 26th Annual Conference 1, 149–153 vol.1 (1996).

3. Yerion, K. A. & Rinehart, J. A. Guidelines for Collaborative Learning in Computer Science. SIGCSE Bull. 27, 29–34 (1995).

4. Streibel, M. J. Queries About Computer Education and Situated Critical Pedagogy. Educ. Technol. 33, 22–26 (1993).

5. Tedre, M., Simon & Malmi, L. Changing aims of computing education: a historical survey. Comput. Sci. Educ. 28, 158–186 (2018).

6. Randall, D. C., Moore, C. & Carvalho, I. S. An international collaboration to promote inquiry-based learning in undergraduate engineering classrooms. Campus-Wide Inf. Syst. (2012). doi:10.1108/10650741211253859

7. Newell, S. Collaborative Learning in Engineering Design. J. Coll. Sci. Teach. 19, 359–362 (1990).

8. Filippi, A. & Agarwal, D. Teachers from Instructors to Designers of Inquiry-Based Science, Technology, Engineering, and Mathematics Education: How Effective Inquiry-Based Science Education Implementation Can Result in Innovative Teachers and Students. Sci. Educ. Int. (2017).

9. Church, G. M., Elowitz, M. B., Smolke, C. D., Voigt, C. A. & Weiss, R. Realizing the potential of synthetic biology. Nat. Rev. Mol. Cell Biol. 15, 289–294 (2014).

10. Way, J. C., Collins, J. J., Keasling, J. D. & Silver, P. A. Integrating Biological Redesign: Where Synthetic Biology Came From and Where It Needs to Go. Cell 157, 151–161 (2014).

11. El Karoui, M., Hoyos-Flight, M. & Fletcher, L. Future Trends in Synthetic Biology—A Report. Front. Bioeng. Biotechnol. 7, 175 (2019).

12. The Future Is Synthetic Biology. Cell (2018). doi:10.1016/j.cell.2018.10.036

13. Gardner, T. S. & Hawkins, K. Synthetic biology: evolution or revolution? A co-founder’s perspective. Curr. Opin. Chem. Biol. 17, 871–877 (2013).

14. Simons, M. The Diversity of Engineering in Synthetic Biology. Nanoethics 14, 71–91 (2020).

15. Alvargonzález, D. Multidisciplinarity, interdisciplinarity, transdisciplinarity, and the sciences. Int. Stud. Philos. Sci. (2011). doi:10.1080/02698595.2011.623366

16. Lawrence, R. J. Deciphering Interdisciplinary and Transdisciplinary Contributions. Transdiscipl. J. Eng. Sci. (2010). doi:10.22545/2010/0003

17. Rapport, D. J. Transdisciplinarity: transcending the disciplines. Trends Ecol. Evol. 12, 289 (1997).

18. Shapira, P., Kwon, S. & Youtie, J. Tracking the emergence of synthetic biology. Scientometrics (2017). doi:10.1007/s11192-017-2452-5

19. Agapakis, C. M. Designing synthetic biology. ACS Synthetic Biology (2014). doi:10.1021/sb4001068

20. Delgado, A. & Porcar, M. Designing de novo: interdisciplinary debates in synthetic biology. Syst. Synth. Biol. 7, 41–50 (2013).

21. Calvert, J. & Martin, P. The role of social scientists in synthetic biology. EMBO Rep. 10, 201–204 (2009).

22. Wang, F. & Zhang, W. Synthetic biology: Recent progress, biosafety and biosecurity concerns, and possible solutions. J. Biosaf. Biosecurity (2019). doi:10.1016/j.jobb.2018.12.003

23. Kuldell, N. Authentic teaching and learning through synthetic biology. Journal of Biological Engineering (2007). doi:10.1186/1754-1611-1-8

24. Stark, J. C. et al. BioBitsTM Bright: A fluorescent synthetic biology education kit. Sci. Adv. (2018). doi:10.1126/sciadv.aat5107

25. Huang, A. et al. BiobitsTM explorer: A modular synthetic biology education kit. Sci. Adv. (2018). doi:10.1126/sciadv.aat5105

26. Stark, J. C. et al. BioBits Health: Classroom Activities Exploring Engineering, Biology, and Human Health with Fluorescent Readouts. ACS Synth. Biol. (2019). doi:10.1021/acssynbio.8b00381

27. Malcolm Campbell, A. et al. pClone: Synthetic biology tool makes promoter research accessible to beginning biology students. CBE Life Sci. Educ. (2014). doi:10.1187/cbe.13-09-0189

28. Anderson, D. A., Jones, R. D., Arkin, A. P. & Weiss, R. Principles of synthetic biology: a MOOC for an emerging field. Synth. Biol. 4, (2019).

29. Mitchell, R., Dori, Y. J. & Kuldell, N. H. Experiential Engineering Through iGEM-An Undergraduate Summer Competition in Synthetic Biology. J. Sci. Educ. Technol. (2011). doi:10.1007/s10956-010-9242-7

30. McLellan, E., MaCqueen, K. M. & Neidig, J. L. Beyond the Qualitative Interview: Data Preparation and Transcription. Field methods (2003). doi:10.1177/1525822X02239573

31. MacQueen, K. M., McLellan, E., Kay, K. & Milstein, B. Codebook development for team-based qualitative analysis. Field methods (1998). doi:10.1177/1525822X980100020301

32. Landis, J. R. & Koch, G. G. The Measurement of Observer Agreement for Categorical Data. Biometrics (1977). doi:10.2307/2529310

33. Lee, E. K., Avgar, A. C., Park, W. W. & Choi, D. The dual effects of task conflict on team creativity: Focusing on the role of team-focused transformational leadership. Int. J. Confl. Manag. (2019). doi:10.1108/IJCMA-02-2018-0025

34. Dimas, I. D., Rebelo, T., Lourenço, P. R. & Rocha, H. A Nonlinear Dynamical System Perspective on Team Learning: The Role of Team Culture and Social Cohesion. in Lecture Notes in Computer Science (including subseries Lecture Notes in Artificial Intelligence and Lecture Notes in Bioinformatics) (2019). doi:10.1007/978-3-030-24302-9_4

35. Vestal, A. & Mesmer-Magnus, J. Interdisciplinarity and Team Innovation: The Role of Team Experiential and Relational Resources. Small Gr. Res. (2020). doi:10.1177/1046496420928405

36. Rodríguez-Sánchez, A. M., Devloo, T., Rico, R., Salanova, M. & Anseel, F. What Makes Creative Teams Tick? Cohesion, Engagement, and Performance Across Creativity Tasks: A Three-Wave Study. Gr. Organ. Manag. (2017). doi:10.1177/1059601116636476

37. Dimas, I., Lourenço, P., Rebelo, T. & Rocha, H. Maximizing Learning Through Cohesion: Contributions From a Nonlinear Approach. Small Gr. Res. 104649642094448 (2020). doi:10.1177/1046496420944488

38. Orwell, G. 1984. (Secker & Warburg, 1949).

39. Hurst, A. & Mostafapour, M. Conflict in Capstone Design Teams: Sources, Management, and the Role of the Instructor. in 4 (2018).

40. De Dreu, C. K. W. & Weingart, L. R. Task versus relationship conflict, team performance, and team member satisfaction: A meta-analysis. J. Appl. Psychol. (2003). doi:10.1037/0021-9010.88.4.741

41. Takai, S. & Esterman, M. Towards a better design team formation - A review of team effectiveness models and possible measurements of design-team inputs, processes, and outputs. in Proceedings of the ASME Design Engineering Technical Conference (2017). doi:10.1115/DETC2017-68091

42. Slimani, S., Ferreira Da Silva, C. & Médini, L. Conflicts Mitigation in Collaborative Design. Int. J. Prod. Res. 44, 1681–1702 (2006).

43. Deci, E. L., Eghrari, H., Patrick, B. C. & Leone, D. R. Facilitating Internalization: The Self-Determination Theory Perspective. J. Pers. (1994). doi:10.1111/j.1467-6494.1994.tb00797.x

44. Deci, E. L. & Ryan, R. M. The Support of Autonomy and the Control of Behavior. J. Pers. Soc. Psychol. (1987). doi:10.1037/0022-3514.53.6.1024

45. Ryan, R. M. & Deci, E. L. Self-determination theory and the facilitation of intrinsic motivation, social development, and well-being. Am. Psychol. (2000). doi:10.1037/0003-066X.55.1.68

46. Jeong, S., McLean, G. N., McLean, L. D., Yoo, S. & Bartlett, K. The moderating role of non-controlling supervision and organizational learning culture on employee creativity: The influences of domain expertise and creative personality. Eur. J. Train. Dev. (2017). doi:10.1108/EJTD-03-2017-0025

47. Kusano, S. M. & Johri, A. Student autonomy: Implications of design-based informal learning experiences in engineering. ASEE Annu. Conf. Expo. Conf. Proc. (2014).

48. Cooper, K. M., Gin, L. E., Barnes, M. E. & Brownell, S. E. An exploratory study of students with depression in undergraduate research experiences. CBE Life Sci. Educ. (2020). doi:10.1187/cbe.19-11-0217

49. Portes, A. The Two Meanings of Social Capital. Sociol. Forum (2000). doi:10.1023/A:1007537902813

50. Kovalchuk, S., Ghali, M., Klassen, M., Reeve, D. & Sacks, R. Transitioning from university to employment in engineering: The role of curricular and co-curricular activities. in ASEE Annual Conference and Exposition, Conference Proceedings (2017). doi:10.18260/1-2--29043

51. Millunchick, J. M. & Zhou, Y. What Affects Student Outcomes More: GPA or participation in co-curricular activities? in ASEE Annual Conference and Exposition, Conference Proceedings 28696 (2020).

52. Krathwohl, D. R. A revision of bloom’s taxonomy: An overview. Theory into Practice (2002). doi:10.1207/s15430421tip4104_2

53. Cartile, A., Marsden, C. & Liscouet-Hanke, S. Teaching and learning design engineering: What we can learn from co-curricular activities. Proc. Can. Eng. Educ. Assoc. (2019). doi:10.24908/pceea.vi0.13756

54. Klassen, M. et al. Engineering: Moving Leadership From the Periphery to the Core of an Intensely Technical Curriculum. New Dir. student Leadersh. (2020). doi:10.1002/yd.20373

55. Tomaska, L. Training biology’s new romantics. EMBO Rep. (2011). doi:10.1038/embor.2011.56

56. Chiel, H. J., McManus, J. M. & Shaw, K. M. From Biology to Mathematical Models and Back: Teaching Modeling to Biology Students, and Biology to Math and Engineering Students. CBE—Life Sci. Educ. 9, 248–265 (2010).

57. Roxå, T. & Mårtensson, K. Microcultures and informal learning: a heuristic guiding analysis of conditions for informal learning in local higher education workplaces. Int. J. Acad. Dev. (2015). doi:10.1080/1360144X.2015.1029929

58. Gregorowius, D. & Deplazes-Zemp, A. Societal impact of synthetic biology: Responsible research and innovation (RRI). Essays Biochem. (2016). doi:10.1042/EBC20160039

59. Betten, A. W., Rerimassie, V., Broerse, J. E. W., Stemerding, D. & Kupper, F. Constructing future scenarios as a tool to foster responsible research and innovation among future synthetic biologists. Life Sci. Soc. Policy (2018). doi:10.1186/s40504-018-0082-1

60. Bammer, G. Should we discipline interdisciplinarity? Palgrave Communications (2017). doi:10.1057/s41599-017-0039-7

61. Bammer, G. et al. Expertise in research integration and implementation for tackling complex problems: when is it needed, where can it be found and how can it be strengthened? Palgrave Commun. (2020). doi:10.1057/s41599-019-0380-0

62. de Lorenzo, V. et al. The power of synthetic biology for bioproduction, remediation and pollution control. EMBO Rep. (2018).

63. Tripp, B. & Shortlidge, E. E. A framework to guide undergraduate education in interdisciplinary science. CBE Life Sci. Educ. (2019). doi:10.1187/cbe.18-11-0226

64. Mazzocchi, F. Scientific research across and beyond disciplines. EMBO Rep. 20, e47682 (2019).

65. Tripp, B., Voronoff, S. A. & Shortlidge, E. E. Crossing boundaries: Steps toward measuring undergraduates’ interdisciplinary science understanding. CBE Life Sci. Educ. (2020). doi:10.1187/cbe.19-09-0168

66. Duschl, R., Maeng, S. & Sezen, A. Learning progressions and teaching sequences: A review and analysis. Studies in Science Education (2011). doi:10.1080/03057267.2011.604476

67. Duncan, R. G. & Hmelo-Silver, C. E. Learning progressions: Aligning curriculum, instruction, and assessment. Journal of Research in Science Teaching (2009). doi:10.1002/tea.20316

68. Duncan, R. G. & Rivet, A. E. Science learning progressions. Science (2013). doi:10.1126/science.1228692

69. Scott, E. E., Wenderoth, M. P. & Doherty, J. H. Learning progressions: An empirically grounded, learner-centered framework to guide biology instruction. CBE Life Sci. Educ. (2019). doi:10.1187/cbe.19-03-0059

70. Kokotsaki, D., Menzies, V. & Wiggins, A. Project-based learning: A review of the literature. Improv. Sch. (2016). doi:10.1177/1365480216659733

71. MacLeod, M. & van der Veen, J. T. Scaffolding interdisciplinary project-based learning: a case study. Eur. J. Eng. Educ. (2020). doi:10.1080/03043797.2019.1646210

72. Sadler, T. D. Situated learning in science education: Socio-scientific issues as contexts for practice. Stud. Sci. Educ. (2009). doi:10.1080/03057260802681839

73. Bloch, M., Lave, J. & Wenger, E. Situated Learning: Legitimate Peripheral Participation. Man (1994). doi:10.2307/2804509

