## Supplementary Figures for "Challenges in Undergraduate Synthetic Biology Training: Insights from a Canadian iGEM Student Perspective"

**Supplementary Figure S1. Basic Team Demographic Information**

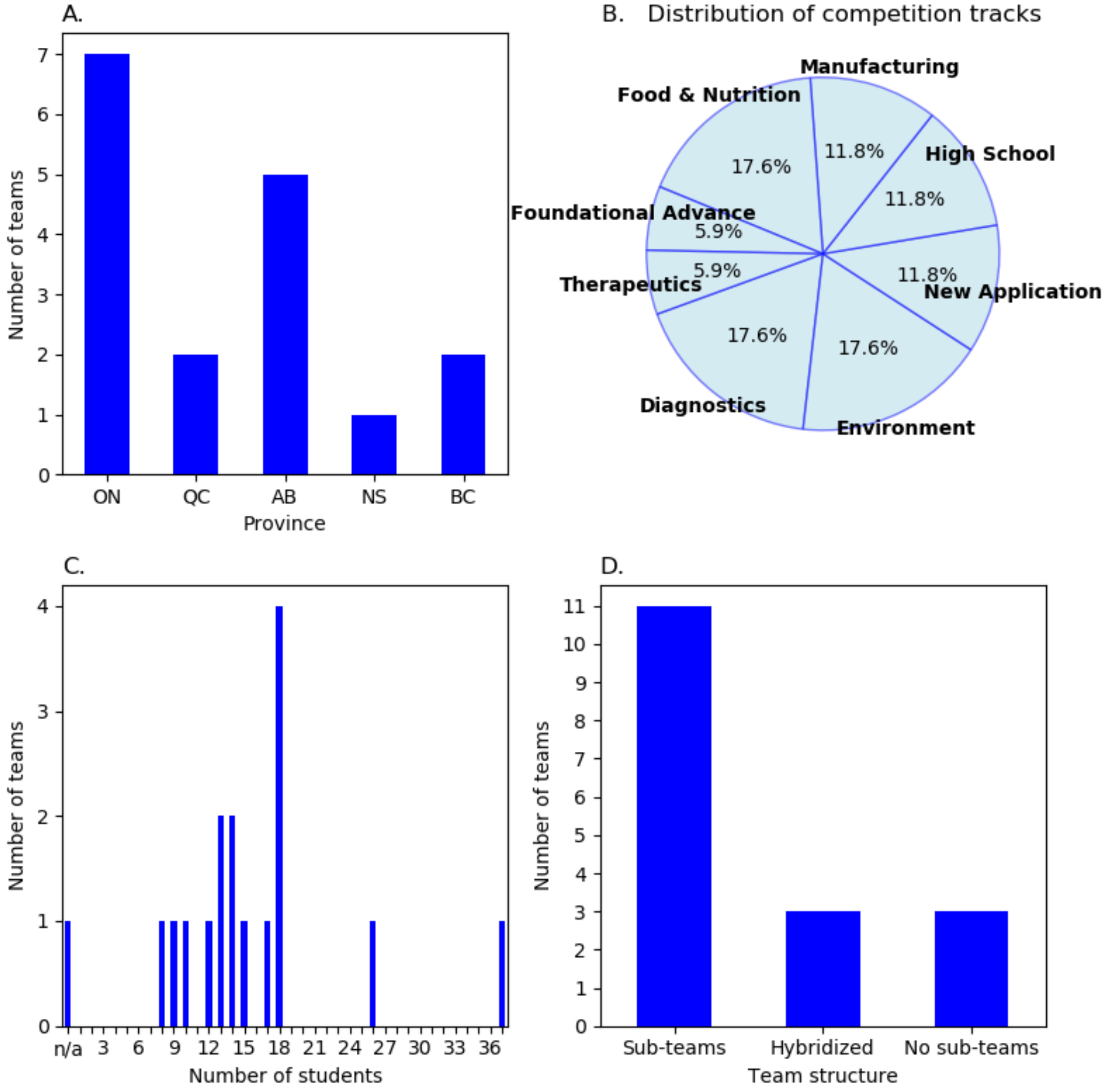


A total of 17 iGEM Canadian teams responded to our e-questionnaire. Twelve of these teams were composed solely of undergraduate students; two were high school teams; three were pre-dominantly comprised of undergraduate students but also included graduate students. Panel A shows the teams are distributed across Ontario, Alberta, British Columbia, Quebec, and Nova Scotia, with the plurality of teams being in Ontario (41%, 7/17) followed closely by Alberta (29%, 5/17). There are no iGEM teams in the other provinces nor territories. The teams reported on their activities in the most recent season in which they competed. iGEM projects are classified into tracks. The distribution of tracks of the surveyed teams is shown in Panel B. The most popular tracks were Diagnostics, Environment, and Food & Nutrition. Conversely, the least popular tracks were Foundational Advance and Therapeutics. The majority of teams (65%, 11/17) have three distinct sub-teams (WL, DL, HP), whereas the remaining 6 teams either have a hybridized team structure in which there are fewer than three sub-teams (18%, 3/17) or have no sub-teams (18%, 3/17). Some teams also reported having a business/ entrepreneurial sub-team.

**Supplementary Figure S2. Team Funding Information**

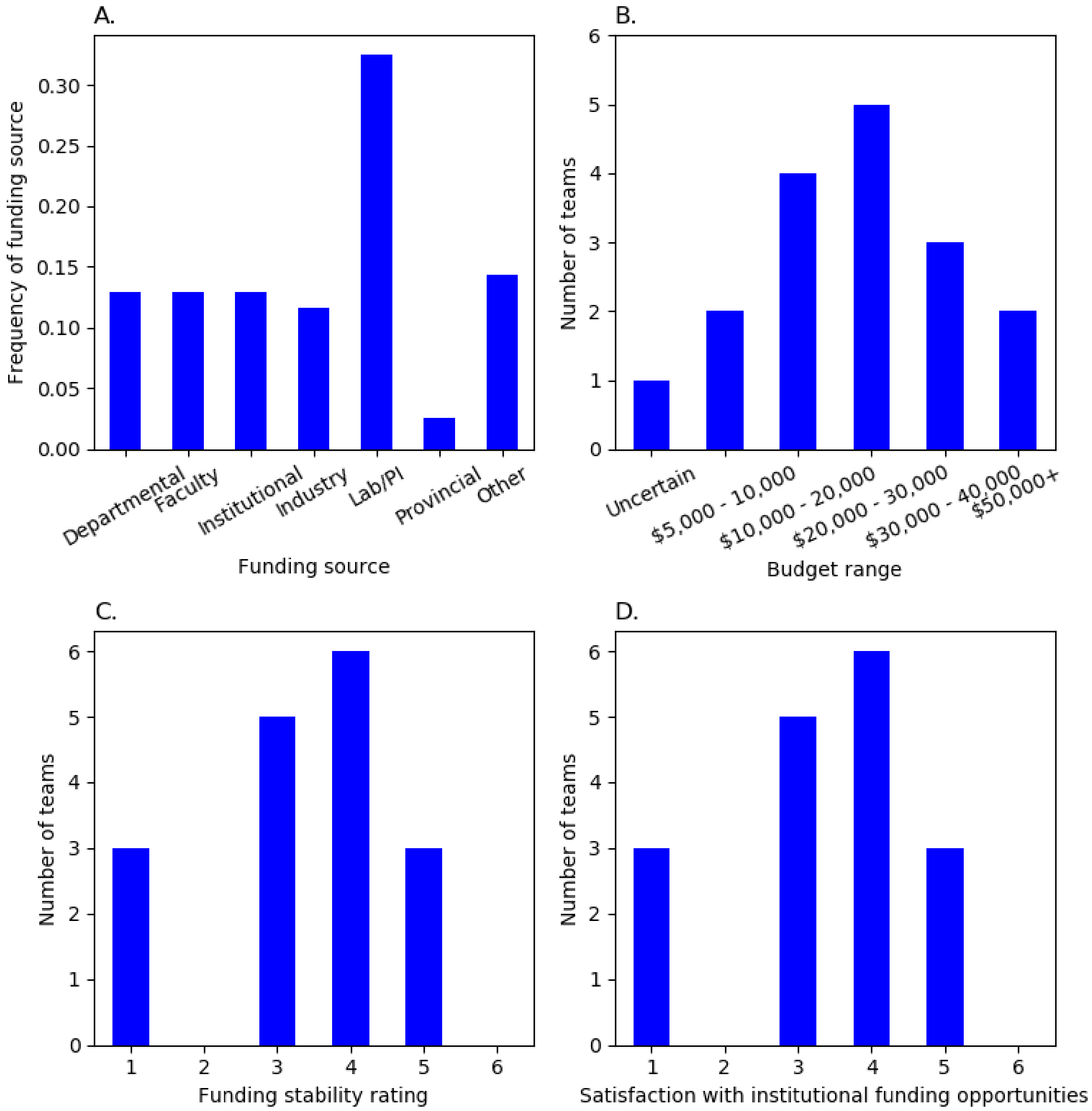


The iGEM teams reported obtaining funding from a variety of sources including departments or faculties at their institution, the institutions themselves, industry (e.g., biotech companies), provincial grants, or from their faculty supervisor (Panel A). Several teams also reported acquiring funding from student unions, team-organized fundraising events, and partnerships with local business. The most prevalent funding source was the team's Lab/PI, followed by "other", which corresponds primarily to funding accrued through team initiatives apart from conventional grant applications or industry partnerships. Provincial funding was uncommon: only two teams, corresponding to an overall frequency of 2.5%. Almost all teams had a total operating budget greater than $10,000 (CAD). Panel B shows the most prevalent budget range was $20,000 – 30,000 (29%, 5/17). Panel C shows approximately half of the teams reported at least moderate funding stability over the past three years, with a rating of 4/6 or greater (53%, 9/17). No teams reported a rating of 6. Nearly half of the teams reported some funding instability. Of these, the most prevalent rating was 3/6 (29%, 5/17). Panel D shows when asked if they were satisfied with the number of funding opportunities available at their institution, approximately half of the teams indicated they were dissatisfied (3/6 or less).

**Supplementary Figure S3. Team Ratings for Support Network Quality**

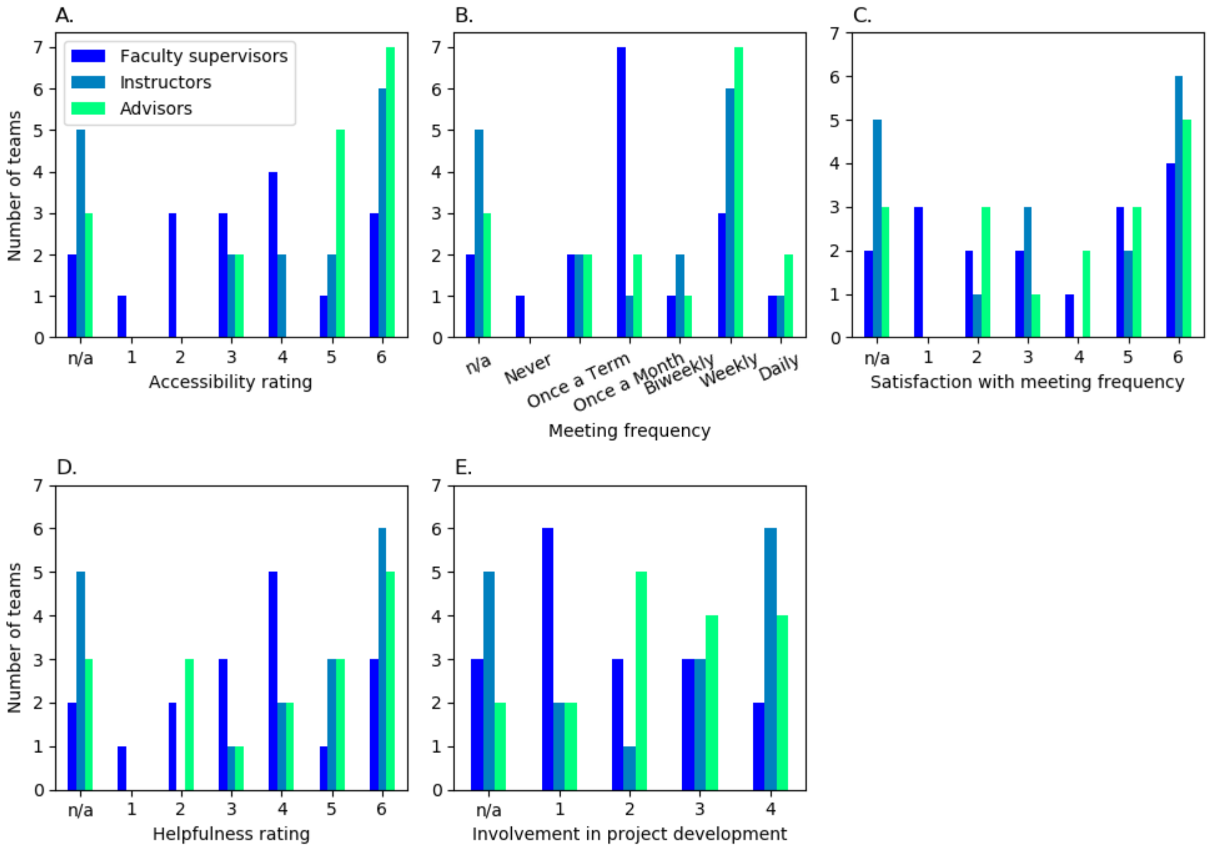


Teams reported on the quality of support provided by their support network (Faculty Supervisors /Instructors /Advisors). At least half of the teams assigned a rating of 4/6 or greater for how accessible their support network was. Advisors were reported to be the most accessible, followed by instructors, then faculty supervisors (Panel A). Panel B shows most teams met with their faculty supervisors at most monthly (71%, 12/17). Conversely, most teams met with their instructors at least biweekly (53%, 9/17) and with their advisors at least weekly (53%, 9/17), with weekly being the most reported frequency in both cases (instructors – 35%, 6/17; advisors – 41%, 7/17). Panel C shows most teams reported at least moderate satisfaction with the frequency of their advisor meetings, assigning a rating of 4/6 or greater (59%, 10/17 ). Approximately half reported a satisfaction rating of 4/6 or greater for the frequency of their meetings with faculty supervisors/instructors (47%, 8/17). Thus, approximately half the teams were relatively dissatisfied with the frequency of meetings with faculty supervisors /instructors. Panel D shows teams reported that their meetings with faculty supervisors /instructors /advisors were helpful to varying degrees. Instructors were most frequently given a helpfulness rating of 4/6 or greater (65%, 11/17), followed by advisors (59%, 10/17), then faculty supervisors (53%, 9/17). While teams tended to rate their meetings with the support network as helpful, they did not report significant involvement of the support network in the development of their projects overall. Panel E shows faculty supervisors and advisors were reported as being the least involved, with most teams assigning an involvement rating of at most 3/6 (71%, 12/17 for faculty supervisors), (65%, 11/17 for advisors). The same statement cannot be made for the instructors. There are many n/a counts for instructors since 5 teams reported having 0 instructors, and for those with instructors the involvement rating for the instructors is relatively spread out with 6 teams assigning a rating between 1 and 3 and 6 teams assigning a rating of 4. No teams assigned an involvement rating of greater than 4 for any of faculty supervisors, instructors, nor advisors.

**Supplementary Figure S4. Prevalence and Demand for Extra-curricular Skills Training**

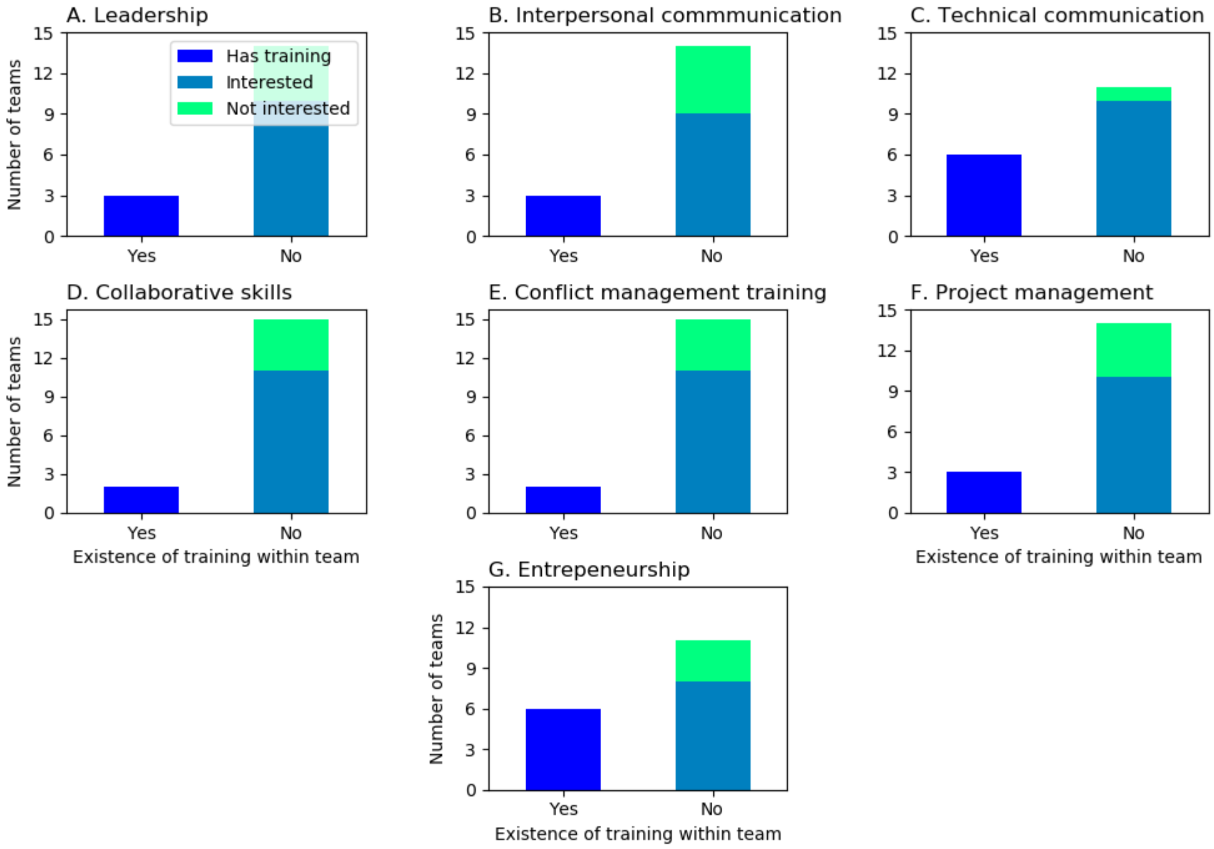


Few teams reported having training available to them for the following skills: leadership, interpersonal communication, technical communication, collaborative skills, conflict management, project management, entrepreneurial training. Moreover, in all cases, teams which reported not having the given type of skills training indicated that they would be interested in this type of training for their members. Of the skills training that teams did receive, technical communication and entrepreneurial skills were the most popular (both – 35%, 6/17), with technical communications the category for which the most interest was shown (90%, 9/10). The least popular were collaborative skills training and conflict management (both – 12%, 2/17).

**Supplementary Figure S5. Timeline of Project Progression**

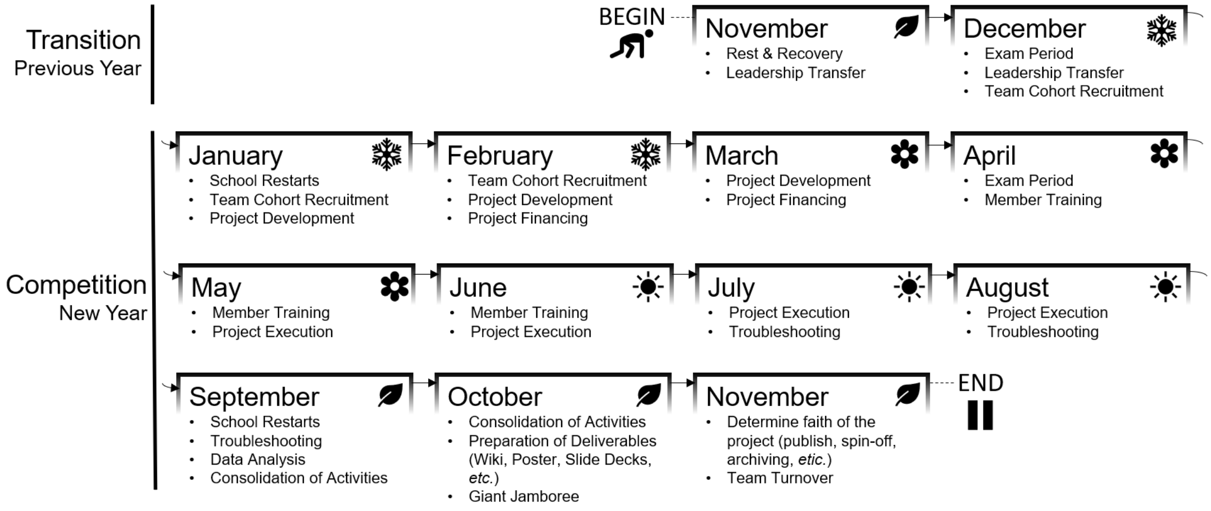


Interview participants were asked open-endedly in Q1 to describe their team's journeys and milestones in their most recently completed competition season. An alignment of these timelines can be collapsed into a canonical progression of Canadian iGEM projects (Figure 1). A new “season” of the iGEM competition began shortly after the conclusion of the previous year’s Jamboree held in late October. Typically, the previous generation transferred leadership responsibilities to the following year's iGEM team cohort in November. Most teams had assembled a complete leadership team for the new iGEM season by the time classes restarted in January of the new year. Additionally, in some cases members of the exiting generation continued with the team by acting as instructors and advisors, facilitating knowledge transfer for logistics, financing, and competition deliverables. Throughout January and February, team leadership  recruited their remaining team cohort, typically to fill roles in three sub-teams: human practices (HP), dry-lab (DL), and wet-lab (WL). HP teams were comprised of students from the arts and humanities (e.g. sociology, philosophy, bioethics) and students from STEM programs who wanted to branch out of their programs’ curricula. DL recruits were comprised of student from mathematics, statistics, physics, computer science, engineering, and quantitative biology programs. WL recruits were drawn from biochemistry, molecular biology, microbiology, biomedical sciences, and engineering programs. Between February and March, most teams brainstormed project ideas. Some teams started project development in January, before completing recruitment of the team cohort, and so were able to begin training by April. However, all teams appear to have slowed down in April due to the examination period.

The bulk of Canadian iGEM activities occurred throughout May to August. Most sub-team training occurred in May and continued through June. Most troubleshooting issues related to WL experimental work typically began in early July (likely because that was the time when the teams’ DNA constructs arrived). In August, project productivity decreased due to students preparing to return to classes. In September, teams reported difficulties with completing the planned summer work and the majority of the work often fell on team leadership as general members focused on re-adjusting to their new class schedules. By late-September, teams began to consolidate their activities into a cohesive story. In October, this story was used to prepare the three competition deliverables that would be evaluated by iGEM judges in the Jamboree: the online project wiki, a conference poster, and the slide deck for an oral presentation. After presenting their work to panels of judges at the Jamboree, awards were announced, and teams again began to re-enter a transition period. Depending on the competition outcomes, teams contemplated a number of options for where to take their projects: academic publishing after additional experiments, embarking on a start-up spin-off of the project, and often times discontinuing the project to allow the next generation to select their own project.
