## Supplementary Materials for "Challenges in Undergraduate Synthetic Biology Training: Insights from a Canadian iGEM Student Perspective"

**Appendix A – Electronic Questionnaire (Questions)**

Part I - Background Information

1. What is your team name?
   Short Answer: _______________

2. What city and province is your team located?
   Short Answer: _______________

3. What type of team are you?
   Radio button options: high school, community lab, collegiate (undergrad), collegiate (grad)

4. What is the name of your institution?
   Short Answer: _______________

5. What track is your team?
   Radio button options: Diagnostics, Energy, Environment, Food & Nutrition, High School, Information Processing, Manufacturing, New Application, Therapeutics, Open, Software

6. How many students competed in your team this year (official roster)?
   Short Answer: _______________

7. How many students in this year's team have competed in previous iGEM teams.
   Short Answer: _______________

8. How many students went to the iGEM competition in Boston?
   Short Answer: _______________

9. How many students do you typically look for in a team?
   *If you do not know this answer, select "Not sure"*
   Radio button ranges: 1-10, 11-20, 21-30, 30+, Not Sure

10. Approximately how many applicants did you receive during your recruitment process for your team?
    *If you do not know this answer, select "Not sure"*
    Radio button ranges: 1-20, 21-50, 51-100, 101-150, 151-200, 201+, Not Sure

11. How many years has an iGEM team existed at your institution? In other words, how old is your iGEM team?
    Radio button range: 1-12

12. Please indicate which years your iGEM team has received a medal:
    Matrix w/ check boxes: 2007 - 2018 (x-axis) and No Medal, Bronze, Silver, Gold (y-axis)

13. Pick the team structure that best matches your team.
    Description: *In this study, we assume there are clear distinctions between wet-lab, dry-lab, and human practices. Wet-lab involves laboratory bench work and molecular biology techniques; Dry-lab involves computational modelling and prediction, as well as computationally intensive bioinformatics, machine learning, and software/hardware design; Human Practices involves stakeholder and public engagement/outreach, educational programs, scientific communications,*

*and policy/sociology-related work.*
Radio button options:
a. Three defined sub-teams (wet-lab, dry-lab, human practices) and a management/administrative/business team
b. Three defined sub-teams (wet-lab, dry-lab, human practices)
c. At-least 1-2 defined sub-teams (*e.g.*, joint wet-/dry-lab).
d. No defined sub-teams
e. Other: _______________

14. What are your team's funding sources (strictly monetary)?
Checkbox options:
a. Individual or Multiple Labs of Principal Investigators (PIs)
b. Departmental
c. Faculty
d. University-wide (*e.g.*, research granting bodies)
e. Industry

15. What is your operating budget this year:
Radio button options:
a. $0 - 5000
b. $5000 - 10'000
c. $10'000 - 20'000
d. $20'000 - 30'000
e. $30'000 - 40'000
f. $40'000 - 50'000
g. $50'000+

16. Our team's funding over the past 3 years have been stable.
Radio button range: Strongly Disagree (1) to Strongly Agree (6)

17. Our institution has had numerous opportunities to apply for funding in the past 3 years.
Radio button range: Strongly Disagree (1) to Strongly Agree (6)

18. Do you have students or student leaders in your team who receive financial compensation for their work on your iGEM project?
If yes, please indicate how many members in the "Other" option.
Radio button options: No, Other ____________

19. If your iGEM team has a website or social media accounts, please provide the URL links below.

Part II - Support Network Access

Your support network is defined as the body of principal investigators (PIs), instructors, and advisors that provide oversight and guidance for your team. The specific definition for these roles can be found here, which we refer to in the following questions: https://2019.igem.org/Competition/Team_Requirements

1. How many PIs, instructors, and advisors does your team have?
Matrix w/ radio buttons, (x-axis) up to 5 and 5+ option; (y-axis) PIs, instructors, advisors

2. To decide on which project to execute for the competition, how much involvement was there from your PIs, instructors, and advisors?

Radio button scale, (x-axis) scale from 1-4, No influence (1), Strong influence (4); (y-axis) PIs, instructors, advisors

3. If you have PIs, instructors, or advisors, how often do you have scheduled meetings with them? Matrix w/ radio buttons, (x-axis) scale from: Never, Once a Term, Once a Month, On a Weekly/Biweekly basis, Daily; (y-axis) PIs, instructors, advisors

4. Your team is satisfied with the frequency which you have scheduled meetings with your support network.
   Matrix w/ radio buttons, (x-axis) scale from 1 (Strongly Disagree) to 6 (Strongly Agree); (y-axis) PIs, instructors, advisors

5. How accessible is help from your support team if you run into an issue in the lab/ on the computer?
   Radio button scale, (x-axis) scale from 1-6, 1 (Not accessible at all) to 6 (Extremely accessible); (y-axis) PIs, instructors, advisors

5. How helpful are your meetings with your support network?
   Matrix w/ radio buttons, (x-axis) scale from 1 - 5, with 5 being “Very Helpful”
   Radio button scale from 1 (Highly unhelpful) to 6 (Highly helpful)

Part III - SynBio Educational Access

1. To what degree do members of your team have access to the following? (The descriptions in brackets refer to which types of teams may select those options).
   Checkbox options:
   a. Course units/ Course themes dedicated to Synthetic Biology
      (High school, Collegiate, Community Lab)
   b. Full Course(s) dedicated to Synthetic Biology
      (High school, Collegiate, Community Lab)
   c. Minors/Specializations/Options with a Synthetic Biology Emphasis
      (Collegiate Only)
   d. Synthetic biology undergraduate programs
      (Collegiate Only)
   e. Synthetic biology graduate programs
      (Collegiate Only)
   f. None of the above.
   g. Other: ______________

2. Are the following opportunities (paid or un-paid) available to students at your institution? Checkbox options: synthetic biology industrial work placements, synthetic biology lab placements, none of the above

3. How does your iGEM team train its members?
   Checkbox options:
   a. No specific training routine. Members are recruited based on self-competency
   b. General learn-on-the-job training as needed
   c. Planned training sessions by student leaders
   d. Planned training sessions by PIs/instructors/advisors
   e. Shadowing individuals with expertise in a field related to the members' tasks
   f. Pre-requisite course to apply to the team that provides sufficient training

    g. Other: ________

4. Does your team or your support network provide training on any of the following for your members? (Yes or No) If "No" for any of the options, please indicate if you think your team would want *and* benefit from training on these topics? (check the "Interested" box if you would be interested)
   Matrix checkbox options:
   Y-axis
    a. Leadership
    b. Teamwork
    c. Interpersonal communications
    d. Technical communications
    e. Collaborative Skills
    f. Conflict management
    g. Project management/Organizational skills
    h. Entrepreneurship/ business skills
   X-axis
    a. Yes, No, Interested

5. Of the following divisions of duties, which one experiences the most difficulty in training members?
    a. Wet-lab
    b. Dry-lab
    c. Human practices

6. What makes it more difficult to train members in that subdivision?
   Long Answer: ______________________________

**Appendix B – Interview Protocol (Questions)**

General

1) Tell us about the story of your team from when you started to when you came back from Boston when you or your members attended the jamboree.
   [*ensure the following questions are sufficiently answered*]
   a) When and how do they recruit and select new members?
   b) How many do they select?
      [*from your judgement if it seems like a lot (+15) or very few (<8), ask why they chose this size*]
   c) Are their members generally from interdisciplinary backgrounds (e.g., biophysics, synthetic biology, bioethics, bio/chemical engineering) which provide training across 2 or more sub-team disciplines, or do they come from more traditional undergrad programs (e.g., pure life sciences, or pure physical sciences, or pure social sciences/ethics/humanities)?
   d) What does their team structure look like?
   e) How is their project formulated?
   f) Does anyone in their support network influence them?

2) What are your plans now for November and December of this year (2019)?
   [*ensure the following question is sufficiently answered*]
   a) Is there a formal way in which they pass on knowledge about the competition AND/OR wet-lab/dry-lab/human practices work to the 2020 team?

3) Please describe the social structure of your team. Are people friends? Do your members participate in social activities outside of your team duties, like having meals together, study groups, seasonal parties, etc.

4) Has your team experienced interpersonal conflict? How have you managed conflict over this past year?
   a) Was there conflict between the iGEM team and the institution?
   b) Does your team have a Code of Conduct, or a Form of Agreement with your institution that establishes clear guidelines?

Support Network

5) The support network is defined as the body of principal investigators (PIs), who are typically professors; and instructors and advisors, who are typically graduate students. We understand that [*describe their response for Part II of the questionnaire*]. Could you describe your support network in more detail?
   [*ensure the following questions are sufficiently answered*]
   a) What is the permanency of the support network?
   b) What type of support do they receive from their support network?
   c) Are there changes they would like to see in their support network?

6) Can you describe what the relationship between your iGEM team and the institution is like, in regards to professors, undergraduate students, and graduate students?
   [*ensure the following questions are sufficiently answered*]
   a) Do professors like iGEM, or are there reservations?

b) Is iGEM something that undergraduate students aspire to be a part of, or is your reputation still building?
c) What do graduate students think about iGEM?

Synbio Education Access

7) How are your iGEM team members trained, both students and student leaders? [*ensure the following questions are sufficiently answered*]
    a) How do you deal with people's gaps in knowledge in sub-team topics other than their own? (*e.g.*, wet-lab member's understanding of biological modelling)
    b) Do your students' programs (*e.g.*, courses, or formal training as a part of their program) prepare them for the work they did on your iGEM team?
        i) Is this answer affected by the program year these participants are in?
    c) How would they make things better? Are there specific resources they need?

8) Please describe any Synthetic Biology courses/Programs that are offered by your institution. If there aren't any, can you imagine what kind of content you would want in the synthetic biology courses?

9) What kind of non-technical skills do you feel like your team members' iGEM experiences have given them? For example: leadership, teamwork, project management, interpersonal communications, etc. Can you specify?

### Appendix C – Code of Conduct Template

[insert iGEM Team Logo]
[iGEM Team Name] Code of Conduct

ARTICLE 1 – PURPOSE AND OBJECTIVES

1. List a purpose/objective of your team at your university (This can include attending the Jamboree, promoting/fostering synthetic biology research, creating opportunities to be involved in multidisciplinary/interdisciplinary project development experiences, etc.)
2. State purpose of this document and general outline of what is to come
3. …

ARTICLE 2 – MEMBERSHIP

1. Outlines eligibility criteria for official members on your iGEM team
2. Can include how one sustains membership, including continued contributions and avoiding professional misconduct
3. ...

ARTICLE 3 – MEMBER ROLES AND RESPONSIBILITIES

It is here where you can define the responsibilities of the different types of members on your team. You can subdivide this into sections if needed, for example administrative vs. research team, sub-team division, etc. Responsibilities should be clearly outlined and defined for each of the different member positions (see below for an example)

The President (1):

- is the only authorized spokesperson of the club
- is one of the designated owners of the club's bank account
- must ensure all the objectives listed in the club code of conduct are followed
- must hold a weekly meeting with the executive committee during their term
- must hold a weekly meeting with the research and development committee during the 10-month research program
- must hold a monthly general meeting during the 10-month research program
- is primarily responsible for formulating administrative and research project based-goals during their term
- is primarily responsible to ensure the completion of all internal and external administrative tasks
- is expected to assume the duties of any member in the executive committee upon his/her/their absence

ARTICLE 4 - ADVISOR ROLES AND RESPONSIBILITIES

Here you can define the roles and responsibilities of your team advisors, in a similar format to what is seen in article 3. Having this section will help ensure that all members of the iGEM team clearly understand the expectations of the advisors, and how they will serve the team. This can include how much help the advisors would provide to the team and to what capacity they would do so. Additionally, it can include how often advisors would meet with the team as a scheduled meeting, or perhaps indicate their availability to be contacted (like office hours).

ARTICLE 5 -SUPERVISOR(S) ROLES AND RESPONSIBILITIES

Here you can define the roles and responsibilities of your team supervisor(s), in a similar format to what is seen in article 3. Having this section will help ensure that all members of the iGEM team clearly understand the expectations of the supervisor(s), and how they will serve the team. This can include how much you would want your supervisor to be involved in the project, schedule meeting times, etc.

ARTICLE 6 - CONFLICT RESOLUTION STRATEGY

In this section you can define the strategies that will be taken when there is internal or external conflict with the team. Internal conflict is defined as problems that arise from within the team itself, while external conflict is a problem that arises involving someone from outside the team. When initially creating this section, it should be discussed with all members of the iGEM team, and reviewed by team advisors and supervisors. The format of this section can be as follows:

Internal Conflict Resolution

- Conflict faced or seen between two individuals will be immediately directed to the lead(s) of the member's respective teams, and the lead(s) will decide to talk/discuss to the involved parties separately or together about the overall situation. From there it would be the lead(s) decision to figure out the next steps in resolving the problem.
- If conflict cannot be resolved with the lead(s), it will be taken up to the advisor(s)/supervisor(s)
- It is the responsibility of the team members to speak up if conflict is seen or heard, to be communicative and not hide problems that may exist

External Conflict Resolution

- Conflict faced or seen between an individual of the team and a third party will be immediately directed to the lead(s) of the member's respective team.

ARTICLE 6 – TERMINATION OF MEMBERSHIP

This is where you would describe how a member will be removed from the team. Make sure to define these guidelines clearly, and you may want to create a process by which a team member's termination may be carried out (for example a general vote by the team, a trial with executive members as the judge, President carrying all power, etc.). Also make sure to include how a team member may defend oneself if put up for termination. Be clear on the consequences should a member termination proceed.

This termination clause is not only limited to the student members, but should also include the advisor(s) and supervisor(s) as well. This is to ensure that all individuals who are involved with the iGEM team, regardless of their role, are held to a high standard and are treated fairly.

ARTICLE 7 – CONSTITUTION AMENDMENTS

As this is an official document for your team to hold your team accountable, it is important to have a section on how changes to this document will commence, so that it cannot be changed at random. It is good to define a standardized process in which revisions can be made, whether during the transition period between iGEM seasons and teams or by other means. It is recommended that the contents of the code of conduct are not changed too greatly during the project year as this may have potential consequences including individuals not working to the standard that they were initially held to. It is advised that major changes to the Constitution during the year are therefore performed with caution.

ARTICLE # – other sections

If you have something specific that pertains to the running of your particular team and has not been aforementioned, then feel free to include more sections as you see fit. Also feel free to rearrange the above articles to the order that would be best for the team. There is no right or wrong answer!

A few other sections that could be included:

- Team policies/guidelines for work conduct
- Determining project for the year (if it is a team decision)
- Who goes to the iGEM competition?
- Who presents at the competition?

ARTICLE AGREEMENT

The final section where all members must sign-off in agreement with this code of conduct (almost like a contract). This would include a section for student members, advisors and supervisors. There is also the option of making different signing sections for each of the respective parties and adding the requirement of having members initial each page (to symbolize acknowledgement of specific clauses), before signing off at the end to finalize the agreement.

### Appendix D – The Support Network/ Institutional Support (SNIS) Scoring Rubric

The Support Network is scored based on the number of criteria they satisfy. There are five criteria: permanency, abundance, availability, quality, championship. To satisfy a criterion in this study, participants only needed to positively comment on the criterion in the interview. We did not evaluate the extent of the positivity expressed for each criterion. Recommendations for how to improve the support network score is detailed in Appendix E.

- Permanency describes the amount of turnover for individuals in the support network during the competition and between multiple competition seasons. Note: not *every* individual in the support network must be permanent, but there must exist a constant individual over multiple competition seasons.
- Abundance describes the number of individuals in the support network and the diversity of individuals in faculty, staff, and graduate positions. It also includes members from the exiting generation that are supporting the team, but are not in the team's roster for that year.
- Availability describes the frequency of the interactions and is related to the DiTO matrix described in Meta-Theme 2. Note: more interactions are not necessarily good for the team. One must consider how much interaction the team wanted with individuals in the support network for the students to work effectively.
- Quality describes the nature of the interactions and is related to DiTO matrix described in Meta-Theme 2. Note: in this context, quality refers to how well individuals in the support network empower students to be comfortably autonomous in their work environment.
- Championship describes individuals in the support network interacting with the host institution in a manner that promotes iGEM.

| Score | Qualifier | Criteria |
|---|---|---|
| 5 | Excellent | All five criteria are checked off. |
| 4 | Good | Four criteria are checked off. |
| 3 | Satisfactory | Three criteria are checked off. |
| 2 | Poor | Two criteria are checked off. |
| 1 | Minimal | One criterion is checked off. |
| 0 | None | No criteria are checked off. |

The Institutional Support is scored based on a single overarching criterion related to the level of engagement the university at-large and faculty express towards the iGEM team. Recommendations for how to improve the support network score is detailed in Appendix F.

| Score | Qualifier | Criteria |
|---|---|---|
| 5 | Excellent | The iGEM competition and the iGEM team's reputation are strongly established at their host institution and their progress is supported/ monitored by the university at-large and faculty leaders. Synthetic biology is fully recognized as an emerging discipline, and there exists a large student community interested in the field. |
| 4 | Good | The iGEM competition and the iGEM team's reputation are established at their host institution and their progress is supported by faculty leaders. Synthetic biology is recognized as an emerging discipline, and there exists a student community interested in the field. |
| 3 | Satisfactory | The iGEM competition and the iGEM team's reputation are growing at their host institution, and some faculty are interested. Synthetic biology is recognized as an emerging discipline among some faculty, and there exists a small student community interested in the field. |
| 2 | Poor | The iGEM competition and the iGEM team have a presence at their host institution, but faculty interest is limited. Synthetic biology might be recognized as an emerging discipline among faculty, and a niche of students are familiar with it. |
| 1 | Minimal | The iGEM competition and the iGEM team have a small presence at their host institution. Synthetic biology may or may not be recognized as an emerging discipline among faculty, and only a small niche of students are familiar with it. |
| 0 | None | The iGEM competition and the iGEM team do not have an established reputation at their host institution. Synthetic biology may or may not be recognized as an emerging discipline among faculty, and is unfamiliar to the student body. |

### Appendix E –Strategic Planning Template for Support Network Engagement

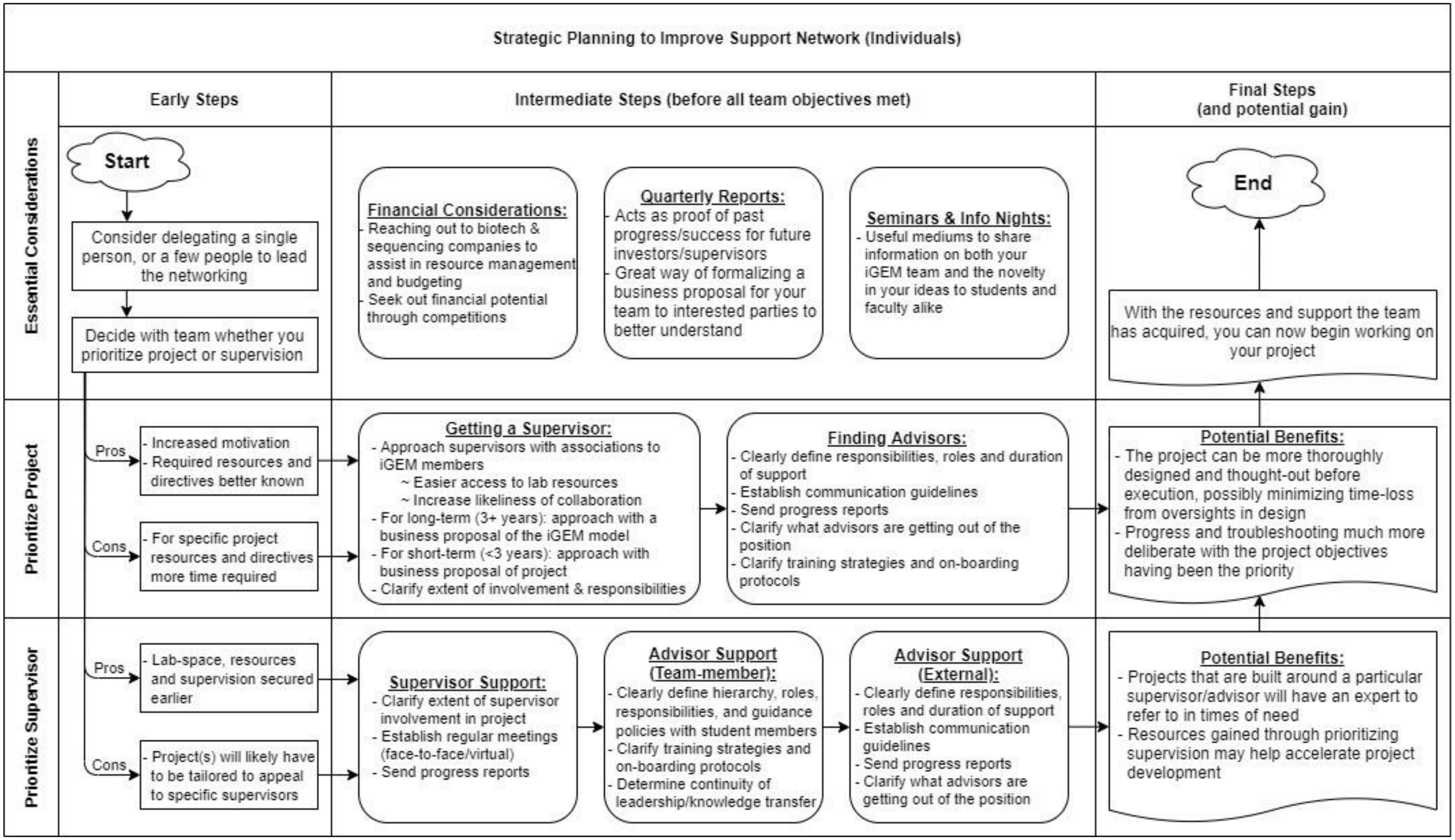

**Appendix F –Strategic Planning Template for Institutional Engagement**

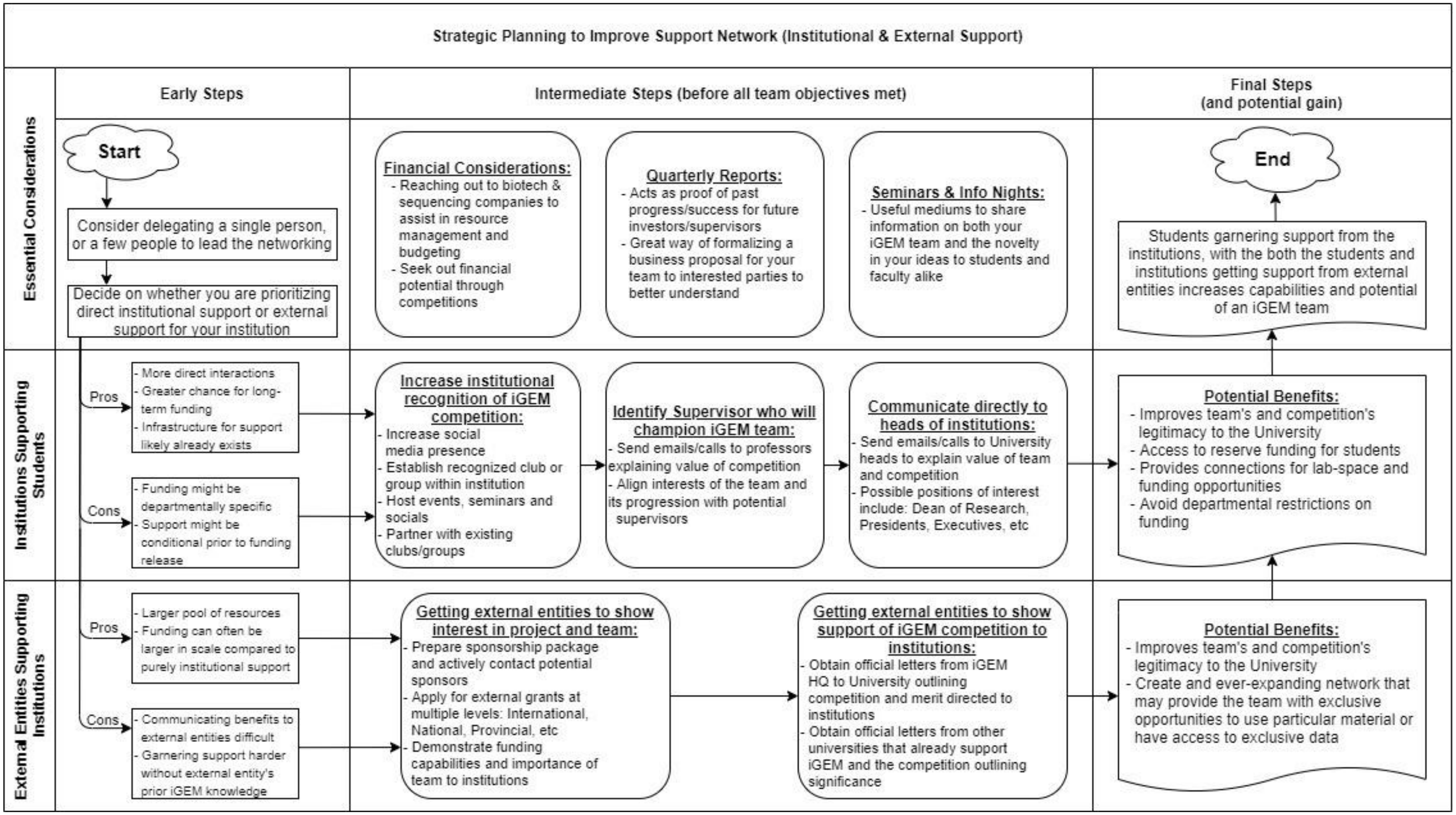

**Appendix G – List of Synthetic Biology or Synthetic Biology related Courses in Canada**
blue = lecture-based, red = team/seminar-based, black = not enough information to determine

| Undergraduate Courses | Course description | Year | Pre-requisites | Course description | Learning objectives |
|---|---|---|---|---|---|
| **British Columbia** | | | | | |
| Victoria- MICR405 (Biotechnology and Synthetic Biology) | Content will cover 7 sections: 1. Cloning, PCR & Sequencing, 2. DNA sequencing and other technologies, 3. DNA and genomic assembly, 4. Elements of genetic circuits, 5. Recombineering and genomic engineering, 6. Making and expressing products, 7. Examples of applications. This course is delivered via a 'flipped classroom' approach. Students are expected to view the lectures (in PDF or audio presentations of PP slides) on the course website before class time. Class time will be used for group learning and independent learning (3hrs a week). | 4 | MICR200A- Introductory microbiology part 1, MICR200B- Introductory microbiology part 2, BIOC300A- General biochemistry part 1, BIOC300B- General biochemistry part 2 | Midterm = 30%, Exam = 35%, Group project = 35% | Students will learn an introduction to how enzymes are used in biotechnology, PCR, cloning, generating cDNA, BACs and YACs, DNA sequencing (including next generation sequencing), Bio bricks and golden gate, genome assembly methods, natural and synthetic promoters, sRNA, CRISPR-based engineering, recombination, RNAi, directed evolution, bioreactors, proteins with un-natural AAs, recombinant vaccines and antibodies, intellectual property related to biotechnology. |
| **Alberta** | | | | | |
| University of Alberta- MICRB315 (Applied Microbiology and Biotechnology) | The synthesis of microbes for commercially used metabolites, drugs, and food enzymes and chemicals. Further processing and modification of the microbes is required for the use of industrial processes. | 3 | MICRB265, this is a second-year course: introduction to microbiology | Instruction through lectures, web-based fermenter simulations, reading assessments, team research, presentations, peer-review exercises. Midterm = 20%, Exam = 35%. Other assessments = 45% | Students will learn about the use of microbes in industry processes. From small-scale in-lab microbiology to large-scale applications, students will learn the process of engineering microbes. They will also learn the interface between microbiology and engineering techniques through steps of action. This may include, isolation and improvement of strains, scaling production, fermentation, purification, integration, and patent rights. Students will also learn about the social issues pertaining to biotechnology applications/ GMOs/ emerging technologies. |
| University of Alberta- BIOCH481 (Design and construction of Synthetic Biological Systems I) | This is the first of 2 synthetic biology classes offered. It offers a theoretical and practical approach to learning the principles of synthetic biology. The topics covered include natural verses the artificial designs of genetic circuits and devices. Students will learn about the experimental aspects of gene and gene network construction as well as their metabolic network design and evaluation. Computer | 3 or 4 | Registered in the faculty of science or engineering with a minimum GPA of 3.3. | Lectures, class discussion, assigned readings, and case studies. | In this introductory class, students will design the basics about protein modelling and design. Students from non-biochemistry backgrounds are encouraged to take the course to amplify the fact that synthetic biology is about the multidisciplinary nature of the field. All information learned will be standardized to accommodate students from different backgrounds. |

| | | | | | |
|---|---|---|---|---|---|
| | modelling, testing, and optimization of proteins will be introduced. | | | | |
| University of Alberta- BIOCH482 (Design and construction of Synthetic Biological Systems II) | This is the second part of the 2 synthetic biology classes offered. This course offers more immediate independence for the students in synthetic biology related projects. It prepares students for participation in the iGEM competition. Team-based problem solving is encouraged. | 3 or 4 | BIOCH481 (Design and construction of Synthetic Biological Systems I) | Design of a mock iGEM project with a presentation and report. | Students will be able to identify a problem in the realm of synthetic biology and clearly devise a plan to solve an aspect of the problem being looked at. Dry lab techniques such as computer modelling to demonstrate feasibility of the design will be explored. The business model and human practices aspect of the project will deem terms under which determine the financial, human, and technological resources to ensure timely completion of the project. An ultimate plan and execution of the plan through a final report and presentation should allow the students to deliver the results to their audience. |
| Victoria- MICR405 (Biotechnology and Synthetic Biology) | Content will cover 7 sections: 1. Cloning, PCR & Sequencing, 2. DNA sequencing and other technologies, 3. DNA and genomic assembly, 4. Elements of genetic circuits, 5. Recombineering and genomic engineering, 6. Making and expressing products, 7. Examples of applications. This course is delivered via a 'flipped classroom' approach. Students are expected to view the lectures (in PDF or audio presentations of PP slides) on the course website before class time. Class time will be used for group learning and independent learning (3hrs a week). | 4 | MICR200A- Introductory microbiology part 1, MICR200B- Introductory microbiology part 2, BIOC300A- General biochemistry part 1, BIOC300B- General biochemistry part 2 | Midterm = 30%, Exam = 35%, Group project = 35% | Students will learn an introduction to how enzymes are used in biotechnology, PCR, cloning, generating cDNA, BACs and YACs, DNA sequencing (including next generation sequencing), BioBricks and golden gate, genome assembly methods, natural and synthetic promoters, sRNA, CRISPR-based engineering, recombination, RNAi, directed evolution, bioreactors, proteins with un-natural AAs, recombinant vaccines and antibodies, intellectual property related to biotechnology. |
| **Manitoba** | | | | | |
| *Brandon- CHEM 15/18.352 (Nucleic acid Biochemistry) ---- Some SynBio* | The class will be delivered via lecture style (3hrs), with no labs. Interactive classroom polls will be conducted based on the required textbook readings and the lecture notes. These classroom polling questions will not be graded; instead, tutorial questions online will be the ones graded. The course will go in depth into the biochemical properties of nucleic acids (nucleotides, DNA, and RNA). The enzymatic biosynthesis of these nucleic acids will be discussed as well as structure and expression of genes. DNA/protein | 3 | CHEM 18:282- Introductory Analytical Chemistry, CHEM 18:271- Organic chemistry II: reactions and synthesis. Or permission from the instructor. | Midterm #1 = 20%, midterm #2 = 20%, problem sets = 10%, Lit review = 20%, exam = 30% | Students will distinguish, recognize, and identify modified nucleotides of RNA. They will be able to design probes and sequence DNA by using the different techniques learned such as PCR, southern, and northern blotting. Explain the theory behind the processing of different RNAs. Explaining how splicing works and how it can contribute to gene diversity. Explain how mutations can affect the organism molecularly and with its fitness. SynBio introduced in the context of 'future directions' of the |

| | | | | | |
|---|---|---|---|---|---|
| | interactions and the function of different RNAs will be explored. Molecular biology techniques will be discussed such as plasmids, PCR, DNA cloning, DNA sequencing, synthetic biology. | | | | molecular techniques learned in the course. |
| **Ontario** | | | | | |
| Queen's- BIOL507 (Biotechnology) | This course will cover topics in recombinant DNA, GMOs, cloning, stem cells, synthetic biology, genetically engineering organisms into the environment, interspecies hybrids, human genome, eugenics. | 4 | BIOL205- Mendelian and Molecular Genetics. Registration in an honours biology program and min. 2.0 GPA or permission from the department. | Seminar = 25%, term paper = 25%, participation and weekly assignments 22.5%, in-class essay = 27.5%, referee evaluation = 5% | The goals for this course include providing a safe and inclusive space for open discussion about controversial topics in biotechnology. Students will learn the background and current research on a variety of biotechnologies. The major goal of the course is to foster an environment for stimulating conversations and critical analysis of public policy and the value of science. Plant biotech will not be covered in detail. |
| Guelph- MBG4240 (Applied molecular genetics in medicine and biotechnology) | The course covers recombinant DNA technologies, DNA analysis including PCR, RNA analysis including northern blotting and in situ hybridization, genome-wide DNA and RNA analysis including microarrays and next generation sequencing, genetic screens including forward and reverse genetics, identification of biomarkers and molecular diagnosis, gene therapy, transgenic and gene-targeted mice, transgenic plants and animals in biotech. | 4 | MBG3040- Molecular biology of the gene | Best 3 out of 4 quizzes = 3 x 18%, (oral presentation = 25%, 4-page critical review of research proposal = 10%, **OR** research proposal = 25%, 2-page reading notes of the research paper = 5%, 2-page evaluation of oral presentation = 5%), participation in discussion = 3%, mini-assignments 5 x 2% but only 4 will be counted. | The goals for the course include being able to focus on the application of advanced molecular techniques and methods used in genetic research, medicine, and biotech industries. Covered topics include DNA and RNA analysis, molecular diagnosis, gene therapy, and transgenesis. Students will learn and gain understanding about modern molecular biology technology through its applications. |
| Waterloo- BIOL349 (Synthetic biology project design) | The course is aimed at including students from multidisciplinary fields to create a project in synthetic biology. The goal is to create a research proposal that can be adopted by the UWaterloo iGEM team. The proposal will require independent work, discussion, collabs between members and supervisors. | 3 | Level at least 3A. | Participation in weekly journal club and discussion sessions = 15%, Written proposal = 15%, poster presentation = 20%, term paper of the design project = 40%, contribution to class discussions = 10% | Active participation in the discussion is necessary as well as in the weekly journal meetings. This will aid students in developing their literacy skills when analyzing the literature. The written proposal will follow a prelim outline of the components of the final project design, compete with figures and references; the poster presentation will be the end component of this. The term paper will be formatted as a final literature paper for the team with |

| | | | | | |
|---|---|---|---|---|---|
| | | | | | methodological detail for the project. Students will learn how to write a detailed report. |
| Western- BCH3392 (Synthetic biology: Principles and Practice) | This course is designed to be an introduction to the principles of synthetic biology. The lectures will revolve around designing biological constructs, DNA synthesis and assembly, editing, delivery, and installation in the destination organisms. The class will consist of lectures and labs/tuts. The course starts off with an intro into synthetic biology followed by DNA assembly methods. An introduction to iGEM is slotted in as 1 entire lecture hour. Multiple guest lecturers are brought in, including a class on, entrepreneurship, and SynBio and ethics. Students will learn how to work with host organisms (E.coli, B.subtilis, and others). DNA transfer between organisms is preformed. How to build synthetic cells is taught as well as sequencing methods and CRISPR. Finally, the building of synthetic organelles is presented. | 3 | BCH2280A- Biochemistry and Molecular biology, BIO2290- Scientific methods in Biology, BIO2382- Cell biology, BIO2581B- Genetics | Midterm = 35%, exam = 30%, ideas presentation = 10%, presentation = 25% | Upon completion of the course the student will be able to retrieve DNA from organisms as well as databases, Use BioBricks for the creation of synthetic biological circuits and be able to explain the positive and negative feedback in the circuits. Become familiar with current projects in academia involving synthetic biology. Be familiar with 10 start-up SynBio companies and understand the science behind their products. Have ethical consideration for building of synthetic organisms/models. |
| Western- BCH4415B (Applications of Synthetic Biology and Chemical Genetics in medicine) | This course has 2 parts: first part is chemical genetics - manipulating the genetic code. Second part is synthetic biology- introduction to recombineering. The first part entails engineering techniques to make proteins; applications in post-translational modifications in bacteria, yeast, and mammalian cells. Also expanding the nucleotide alphabet with an overview of non-canonical DNA and RNA bases with possible applications. The second part includes tools for genome-wide editing: multiplex genome engineering and phage assisted continuous evolution. Also, applications of designer organisms for biofuels, biocontainment, and de-extinction. | 4 | Either (Biochemistry 3381A and Biochemistry 3382A)- Biological macromolecules, Biochemical regulation **OR** (Biochemistry 2280A and one of Chemistry 3393A/B or Chemistry 4493A/B)- Biochemistry and molecular biology, medicinal chemistry, Chemistry of biological macromolecules | Presentations = 5%, take home test = 30%, in-class quizzes = 15%, 'News and views' article = 20%, final take home exam = 30% | Students will have gained understanding of how biosynthetic gene clusters are organized to produce secondary metabolites in bacteria and fungi. Methods to engineer microbial BCGs to produce known metabolites and screen for novel ones. Screening for microbes and enzymes that can degrade secondary metabolites. |
| Brock University- BIOL4P20 (Synthetic Biology) | Topics in synthetic biology including current and emerging technologies with a focus on engineering advances in biology. This course will teach and give students the tools in order to use methods in synthetic | 4 | One of BTEC 3P50 (Molecular Genetics), BCHM 3P01 (Metabolic Biochemistry), BIOL 3P51 (Genetics) | Individual research paper = 30%, Group research proposal/presentation | Students will learn critical methods in analyzing papers and giving paper critiques. Presentation skills will also be attained and improved on because each student will present an overview of a |

| | | | | | |
|---|---|---|---|---|---|
| | biology in topics such as stem cells, CRISPR/Cas9 genome editing, optogenetic control of cellular communication, engineering the immune response, programmable DNA-based materials, and engineered biosensors. | | | = 40%, class participation = 30% | paper using only the relevant data from the papers. Answering insightful questions, understanding of the paper, and the clarity and flow of the presentation will be looked at and judged upon. Students will learn to contribute meaningful and highlighted information from the assigned readings and be able to speak about them in a class discussion and debate. |
| Carleton University-BIOC4008 (Computational systems Biology) | Lecture is 1.5 hours a week, with workshop 1.5 hours a week. This class targets modelling and simulation of metabolic and regulatory networks which an emphasis in understanding the dynamic cellular systems. Applications in biotechnology such as synthetic biology, metabolic engineering, and drug discovery are learned upon. The workshop portion includes an experimental learning activity in the course. | N/A | BIOC3101 (General Biochemistry) | N/A | N/A |
| Carleton University-BIOC4204 (Protein Biotechnology) | Advanced lecture, discussion, and seminar course with lecture 2 hours per week and workshop 2 hours a week. The theory behind protein and enzyme engineering is looked upon as well as their development and the current techniques being used. Topics discussed in biotechnology include biotechnology, nanotechnology, and new frontiers in basic and applied research. | N/A | BIOC3101 (General Biochemistry), BIOC3202 (Biophysical techniques and applications), or consent from the institute. | N/A | N/A |
| Ryerson University-BMS750 (Systems Biology) | This course is offered with 3 hours lecture and 1.5 hours of lab time per week. It focuses on the integration of complex biological data and non-reductionist approaches to studying living organisms. Topics include, researching systems theory and stochasticity in biological systems, emergent behaviours, the advent of high-throughput biological experimental techniques and the use of modelling to understand biological processes. Also, this course examines epigenetic systems, lipidomic, metabolomics, synthetic biology, integrative cellular signalling networks and computational modelling of cellular | N/A | BLG411 (Cell Biology II), BLG307 (Molecular Biology) | N/A | N/A |

| | | | | | |
|---|---|---|---|---|---|
| | systems. The labs will complement the lectures. | | | | |
| *University of Ottawa-BCH4172 (Topics on Biotechnology)* | This course is taught via lectures and group discussion. Guest speakers are also invited and introduced to describe various advance sin biotechnology. A lecture and seminar course on the application of molecular biology to the field of biotechnology. Topics to be discussed include, gene cloning, expression systems, use of vectors for gene cloning in prokaryotes and eukaryotes and the production of proteins in heterologous hosts. | 4 | BCH3170 (BIOCHEMISTRY), BIO3170 (Molecular Biology) | N/A | N/A |
| University of Toronto-HPS346 (Modifying and Optimizing Life: on the Peculiar Alliance between AI, Biology, and Engineering) | This course integrates the sociocultural and technological conjuncture that has brought multidisciplinary programs together such as computer science, biology, and engineering. These disciplines should be seen as alliances in the advancement of biotechnology. Additionally, methods to see how AI, synthetic biology, and biotechnology are related/associated, and the debates and ethics that come from associations all these disciplines are discussed. Topics include geoengineering, bioremediation, GMO, and robotic insects and the use of expert systems and machine learning to optimize synthetic biology. The ethics behind CRISPR babies, flourishing and marketing of precision and personalized medicine and immunotherapy. | 3 | 4.0 credits | N/A | N/A |
| University of Toronto-CSB490 (Team-based learning: Current topics in Cell and Molecular Biology) | In this course, students will work in teams of 5 people, or based on class size. There are very minimal lectures in order to focus on self-directed learning and research in a team environment. A particular focus on protein-protein interactions and protein biochemistry is built upon, with the overall theme for the course being 'Regulatory Mechanisms in Plants'. Through established data in a repertoire of protein-protein interactions, analysis of interrogating biological systems with synergistic approaches can be conducted. | 4 | BIO260H1/HMB265H1 (Genetics), CSB330H1 (Techniques in Molecular and Cell Biology) OR CSB349H1 (Eukaryotic Gene Expression) OR CSB352H1 (Bioinformatic Methods) | **Individual Work (Total of 61%)**<br>20% Individual Multiple Choice (M/C) (5% each)<br>4% Individual Participation in 4 Paper Discussions (1% each)<br>26% Individual annotated bibliography<br>6% Self and peer- | Students will learn how to be an involved member of a team, as well as learning how to be dependent of team members for the ultimate project to be successful. Learning how to read and analyze a scientific paper, as well as pulling the relevant information from the paper to present to teammates in an efficient manner. Students will also learn how to write a grant proposal, using other student's feedback in order to give a brief overview of the rationale, |

| | | | | | |
|---|---|---|---|---|---|
| | Through analysis of the data, metabolic regulation and the role of cell signalling and regulatory systems in plants can be discovered. | | | assessment Metrics—done as individuals—i.e. every student evaluates self and peers<br>2% Presentation to the Panel (final oral presentation of proposal, individual component)<br>3% Individual Discussion Participation for 3 proposals (1% each)<br>**Team Work (Total of 39%)**<br>4% Team M/C (1% each)<br>4% Team Presentation of 1 paper topic and Preparation of paper questions for 1 paper (worth 2% each)<br>2% Team Presentation of draft Rationale and Objectives<br>2% Team Presentation to the Panel (final oral presentation of proposal—group presentation as a whole)<br>2% Team Panel Leadership (2%)<br>5% Team log book (draft worth 2% and final log book worth 3%)<br>20% Team Final Proposal | objectives, and methodology presented in class. |
| Concordia-BIOL 512/BIOL 482 Functional Genomics | This course focuses on the functional analysis of expressed genes and their products. Course content includes the construction and screening of normalized cDNA libraries,<br>analysis of expressed sequence tags (ESTs), | Undergrad and Graduate | BIOL 367 (Molecular Biology) | Online quizzes 2 %<br>Class participation, reading assignment, discussion forum 8 %<br>Oral presentation 16 % | Online quizzes: Complete three online quizzes on Moodle. One quiz is to test your knowledge on plagiarism and citation protocols. The second one is to test your knowledge on techniques in molecular biology and on class materials. |

| | | | | | |
|---|---|---|---|---|---|
| | functional analysis by gene knock-outs, localization of<br>gene products by gene knock-ins, transcription profiling, systematic identification of proteins, and<br>functional analysis of proteins by detection of protein-protein interactions. | | | Assignments 12 % x 2<br>Midterm exam 10 %<br>Final exam 40 % | The third one is to test your knowledge on class materials. Reading assignments: Read the assigned article. Participate in pre-class discussion. Participate in the class discussions. Answer questions regarding the article during the class. Oral presentation: Select and read a latest article (less than three years) on functional genomics. Describe the major finding of the article as oral group presentation. Assignments You will study and characterize a mouse gene using publicly available expression and phenotype databases. Direction will be posted on Moodle. |
| UQTR- TSB1001-00 Biological Engineering | Introducing Recombinant DNA Technology and Methods Underlying Current Rise<br>of the biotechnology sector with reference to the applications developed.<br>Reminder of molecular biology.<br>Mechanism of translation. Enzymatic tools and vectors of<br>cloning. Methods of analysis and detection associated with bioengineering. The reaction of<br>chain polymerization and its applications.<br>Site-directed mutagenesis, transgenesis, and engineering<br>proteins. | 2 | BCM1002 Biochimie II | Research plan 5%<br>Midterm 25%<br>Assignment 20%<br>Oral presentation 25%<br>Final 25% | Acquire concepts relating to nucleic acids, the essential concepts concerning structure and expression of genomes as well as the main techniques of analysis and<br>manipulation of genes from bacterial systems to eukaryotic organisms;<br>Understand site-directed mutagenesis and its applications in the construction of organisms<br>transgenic and the production of specific proteins;<br>Understand the main approaches used in the biotechnology industry and the impact that<br>this one on us. |
| University of Sherbrooke-GNT512-Biomolecular engineering | For several millennia, understanding natural mechanisms has enabled humanity to better<br>control her environment and maximize the benefits derive from it. This understanding first manifested itself in the development of agriculture, made possible by the domestication<br>of plant and animal species. It was then applied to selective breeding, allowing the development in several of these species of characteristics appropriate to different needs<br>(better performance, better resistance to parasites or the elements, etc.). Thanks to | Undergrad | GNT310 - Génétique et biologie moléculaire | Midterm (30%)<br>Final Exam (40%)<br>Assignment 1 (7%)<br>Assignment 2 (8%)<br>Assignment 3 (15%) | 1- Understand the basic concepts that make possible the manipulation of genetic material (structure<br>DNA, gene expression mechanisms, protein production).<br>2- Know how to use the main tools and techniques making possible the manipulation of genetic material (enzymes used to analyze, manipulate and modify DNA; means of introducing DNA<br>modified in a prokaryotic or eukaryotic cell; systems to ensure gene expression modified).<br>3- Know how to plan the expression of |

| | | | | | |
|---|---|---|---|---|---|
| | biomolecular engineering, one can now aim to develop such advantageous traits by modifying directly the genetic heritage of target species, by avoiding the vagaries of a genetic crosses. The Biomolecular Engineering course seeks to familiarize students with the theoretical and practical bases of the manipulation of genetic material in bacteria, yeasts, plants and animals, as well as with the many applications made possible by these manipulations. | | | | recombinant proteins as well as their purification.<br>4- Be able to cite and explain examples of applications of biomolecular engineering in industry pharmaceutical, agricultural industry, bioremediation and the medical field. |
| ULaval-BCM-2700-Laboratories on Molecular Biology and Genetic engineering | Molecular biology is and will remain an essential tool for years to come. It is therefore imperative to know the theoretical and practical bases for access to positions in very varied fields, such as agriculture, medical field, education, research, or even forensics.<br>Molecular biology is gaining more and more importance in the life sciences. In agriculture, one cannot pass in silence GMOs, resulting directly from molecular biology (gene manipulation). Besides this aspect controversial, for the moment we can also appreciate the progress in the medical field so much to obtain much more precise, reliable and above all faster diagnoses, reducing the stressful time for the patient while waiting a response for a diagnosis (e.g. cancer). We can also predict that some cancer treatments, which are currently under study or in development in research laboratories, with approaches involving viruses, will emerge in the reasonable future. Forensics Now Uses DNA Evidence of crime scenes, helping to resolve contentious cases, or without evidence until now. She even helped people unjustly accused and imprisoned to prove | Undergrad | BCM-1700-Molecular biology | Bioinformatic exercise Individual 4 %<br>DNA extraction Team 2 %<br>Quiz 1 individual 6 %<br>Cloning team 5 %<br>DNA Extraction Team 3 %<br>RT-PCR team 3 %<br>qPCR DNA Team 3 %<br>Quiz 2 individual 6 %<br>Protein purification Team 2 %<br>Final Individual 9 %<br>Immunodetection team 2 %<br>Laboratory notes individual 10 %<br>Scientific paper team 25 %<br>Oral presentation team 18 %<br>Participation in question period Individual 2 % | Perform DNA study techniques: extraction, purification, mapping, PCR, quantifications;<br>Perform RNA study techniques: purification, reverse transcription (RT-PCR), quantification;<br>Perform protein study techniques: purification, SDS-PAGE gel, solubility, Western;<br>Perform certain genetic transformation techniques;<br>Interpret and analyze the results obtained in the light of knowledge of the operating mechanisms cellular, genetic |

| | | | | | |
|---|---|---|---|---|---|
| | their innocence and thus correct regrettable errors.<br>Regardless of the model organism used, the majority of techniques used in molecular biology laboratories are similar. This course aims to teach the most common methods of molecular biology. | | | | |
| ULaval-BCM-3010-Laboratories on synthetic biology | Through guided exercises, this practical work course familiarizes the student with the basic tools and techniques of molecular genetics and shows him advanced techniques using model organisms. Thus, the student will have to design and carry out a synthetic biology project using the yeast S. cerevisiae as a model organism. At the end of the course, he will be able to develop strategies that will allow him to address the problems of molecular genetics modern. He will then be able to use the techniques of cloning, PCR, gene deletion and insertion and site-directed mutagenesis in vitro. | Undergrad | N/A | Lab report 1 10 %<br>Midterm 25 %<br>Lab note 1 10 %<br>Final 25 %<br>Lab note 2 10 %<br>Lab report 2 15 %<br>Attitude 5 % | 1. The first objective of this laboratory is to familiarize the student with the tools and techniques of basis of genetic engineering.<br>2. The second objective is to allow the student to use his knowledge to design and carry out a project of synthetic biology involving yeast as a model organism.<br>3. The third objective is to enable the student to make a critical synthesis of the results obtained and to analyze them. present in the form of reports respecting the standards of scientific communication. |
| UdM-GCH8650-Biochemical Engineering | Biotechnological production processes, molecules, cells, and tissues. Bacteria, yeasts, fungi, plant, and animal cells. Enzymatic and nutritional kinetics, metabolic pathways, and metabolic engineering. Genetic modifications. Enzymatic and biological reactors. Kinetics of cell growth and production of metabolites including recombinant proteins. Characterization, design, and choice of bioreactors. Sterilization and heat transfer. Theory and practice of mass transfer and scaling of bioreactors. Theory and practice of genetic transformation of cells and culture into a bioreactor of cells and tissues. Product recovery | Undergrad | GCH2105-Biological Engineering | N/A | N/A |
| UdM-GCH2105-Biological Engineering | Engineering of living systems involving biomolecules, biological catalysts, and cells. Fundamental bases and applications related to engineering. Structure and role of the different cellular components. Types of cells and biological catalysts. Examples of bioprocesses involving different types of | Undergrad | GCH2105-Biological Engineering | N/A | N/A |

| | | | | | |
|---|---|---|---|---|---|
| | cells. Cellular nutrition and growth. Aseptic conditions. Bioreactor operating conditions. Proteins: structures and analyzes. Enzymatic kinetics. Study of cell metabolism pathways and metabolic regulation. Mathematical models for problem solving. Understanding of the basics of genetic engineering. Case studies in the fields of production biotechnologies, environmental engineering, biopharmaceutical engineering, biomedical engineering, and agri-food. | | | | |
| **Graduate:** | | | | | |
| **British Columbia** | | | | | |
| Alberta (GRAD)-AFNS522 (Advanced Biocatalytics) | This course provides an approach to whole cell fermentation systems through agricultural sciences using enzyme-based approaches. | Graduate | At least 3 microbiology classes, or have consent form the prof. | Instruction through lectures, peer discussion and debate, guest speakers, and group activities. Midterm = 30%, Exam = 30%, 4-5 assessments = 20%, major project = 20% | Working in small teams will develop the student's sensed of leadership and cooperative learning in order to foster an environment where conflict resolution can be learnt. Independence for the students to organize their own activities and manage their time outside of class-time will be expected. The course will teach the students the theory behind approaches used in bio-industrial food industries. |
| Calgary- CPSC607 (Biological Computation) | Content covered includes cellular automata, evolutionary computing algorithms, swarm optimization, swarm intelligence systems, immune system computing, membrane computing. | Graduate | Consent from the department. | In class presentations = 20%, project implementation = 20%, final project report = 30%, final project presentations = 10%, final oral exam = 20%. NO formal final exam. | To be able to model and examine biological networks. |
| University of Calgary-Medical Science 605 (Information storage and processing in Biological Systems) | This course deals with examining complex biological systems as well as concepts and fundamentals of biological solutions to information storage and processing. Topics of study include modelling and computer simulation of biological systems, information storage in biological molecules, genetic networks, hierarchical organization of biological information processing in signal transduction, development, evolution, biological control systems. | Graduate | Consent from the faculty | N/A | N/A |

| | | | | | |
|---|---|---|---|---|---|
| **Ontario** | | | | | |
| Guelph (GRAD)-BIOT6500 (Molecular biotechnology) | The course will be delivered via modules. Each module will have lecture components and hands-on practical components. Module 1= Applications in Biotechnology including omics and applications, metabolite profile in agriculture, medicinal plant metabolics. Module 2= Tools in biotechnology 1 including DNA technology, chromatography technology, and mass spectrometry. Module 3= Biotechnology and the life science industry including modeling human disease, screening for pre-clinical drug development. Module 4= DNA technology, imaging technology, screening technology. | Graduate | N/A | Class presentations for modules 1&3 = 2 x 25%, class participation = 10%, written data reports for modules 1&2 = 2 x 10%, written data report module 4 = 20%. | Learning outcomes will include demonstrating advanced critical analysis for the scientific literature in order to find a specific research goal. Be able to communicate efficiently through written reports. Understand the global context of biotech and identify its applications. Communicate through written and oral forms. |
| University of Toronto-PHM1140 (Principles of Synthetic Biology) | Using didactic teaching (17 hours) and practical classes (13 hours), in this course, synthetic biology will be explored in relevance to application in the pharmaceutical sciences and beyond; such as in topics that can be applied to student's own research. Additionally, topics covered include genetic circuits and metabolic pathways to construct new smaller molecular drugs and novel protein/RNA based therapeutics. This class is designed as an introduction to synthetic biology, assuming students have no prior knowledge about the field. Lectures are focused on synthetic biology and technologies that are driving the field, along with practical theory on design of genetically encoded tools for the application of human health. | Graduate | N/A | N/A | N/A |
| **Quebec** | | | | | |
| Concordia University-Biol 515: Biotechnology and Genomics Laboratory | Biol 515 is a lab course designed to provide hands-on experience with several genomic/biotechnology approaches and provides the opportunity to apply some of the theoretical knowledge obtained in first term course of the Diploma Program in Biotechnology and Genomics. The lab is composed of 2 main modules, which take several weeks each. There will be some overlap between modules during certain | Graduate | N/A | Basic lab skills (prelab 1): 5 marks<br>Report on Standard lab exercises and PCR: 5 marks<br>Lab report 1: 15 marks<br>Lab report 2: 25 marks<br>Assignment: 5 marks<br>Quizzes: 5 marks<br>Final exam: 30 marks | N/A |

| | | | | | |
|---|---|---|---|---|---|
| | weeks. The modules involve qRT-PCR, affinity purification of TAP-tagged proteins, mass spectrometry and co-immunoprecipitation. Although it normally takes several trials in order to become independently competent with certain experimental techniques, the lab course provides “hands-on” experience with a diversity of molecular and biochemical approaches at the genome-wide level, which in turn contributes a solid base for future work in these areas. | | | General Lab Performance: 5 marks<br>Tutorial participation: 5 marks | |
| University of Sherbrooke-GNT 610-Advanced genetics | Continue training in genetics for students who already have a good foundation in this science. | Graduate | BCL102 & (GNT302 / 304 / 310) | Quizzes: 18 %<br>Assignment 10 %<br>Midterm : 30 %<br>Final : 42 % | N/A |

**Appendix H – Learning progression for genetic circuits, adapted from Scott *et al.* (2019)[69]**

| Learning Objective (Program-level): Students can design genetic circuits for research and industrial purposes | | | |
|---|---|---|---|
| Level | Difficulty | Learning Goals (Course-level) | Course Progression |
| Upper Anchor | Sophisticated scientific ideas | (U2) Students understand how to assemble and model different genetic circuit elements to create biosensors, perform biocomputations, and study naturally existing circuit motifs. | Advanced Genetic Circuit Design (4th Year) |
| | | (U1) Students understand the architecture of the Collins' Toggle switch, repressilators, and oscillators; and are capable of modelling their behavior. | Fundamentals of Genetic Circuit Design (4th Year) |
| Intervening Levels | Emerging scientific ideas | (I6) Students understand how to apply differential equations to modelling transcriptional, translational, and metabolic processes. | Mathematical Modelling of Biological Systems III (3rd Year) |
| | | (I5) Students understand that transcriptional proteins act on each other in systems resembling logic gates. | Introduction to Synthetic Biology (3rd Year) |
| | | (I4) Students understand genes encoding transcriptional proteins can be recombinantly expressed using inducible promoters. | Molecular Biology III (3rd Year) |
| | | (I3) Students learn how to build and analyze models for diverse complex biological phenomena. | Mathematical Modelling of Biological Systems II (2nd Year) |
| | | (I2) Students understand the expression of these transcription-related biomolecules are from other genes, and there is a shared interconnectedness. | Molecular Biology II (2nd Year) |
| | | (I1) Students understand transcription is orchestrated by an array of DNA sequence elements, DNA-binding proteins, and other biomolecules. | Molecular Biology I (2nd Year) |
| Lower Anchor | Initial, naïve ideas | (L2) Students understand transcription in the context of central dogma and cell function. | Introduction to Biochemistry & Molecular Biology (1st Year) |
| | | (L1) Students understand calculus and differential equations can be used to model biological phenomena. | Mathematical Modelling of Biological Systems I (1st Year) |

Beginning in the 1st year of undergraduate studies, students would begin with a quantitative course (L1) and an introductory biology course (L2) to begin constructing basic mental models of synthetic biology from the DL and WL perspective. As they progress through a SynBio undergraduate program, they would construct more complex mental models grounded in molecular biology (I1/I2), molecular cloning (I4), applied biochemistry (I5), and model simulations and analysis (I3/I6). Finally, students would engage in genetic circuit design at higher levels of Bloom's Taxonomy of Learning where they apply their modelling knowledge to analyze the behavior of well-studied genetic circuits (U1), then synthesize their own genetics circuits that they can evaluate using more sophisticated techniques (U2). The overarching learning objective of training students to be able to work with genetic circuits computationally and experimentally, provide instructors with a clear idea of how to design instructional methods and assessment instruments towards a connected series of SynBio courses. Depending on faculty and institutional interests, other SynBio LPs could include operating bioreactors, creating bioinformatics workflows, and developing microfluidic/ cell-free biosensors, among others.

**Appendix I – Study Limitations**

The study design reported here possesses some limitations. The lack of specificity of the single-item measures in the e-questionnaire, as well as some shallow interview questions, led participants to comment on constructs and not the constructs' operationalization. For example, we broadly asked about conflict in teams during the interview and received a wide range of responses from interviewees discussing *only* relational conflict, task conflict, or both. Nonetheless, the goal of this study was to explore the landscape and use a grounded theory approach to generate new ideas to describe the state of Canadian synthetic biology training. Future research should aim to explore more nuanced problems identified within the Canadian landscape described in this report by asking more precise questions about specific operationalizations of success, conflict, autonomy, *etc.*

Because we did not have the resources to survey multiple members on a single team to have a more representative response on behalf of the team, we were constrained to asking only one or two student leaders in the group. Many of these student leaders were the presidents and project co-ordinators of the team (*viz.* the primary contact), so theoretically they possess a representative view of the team since they likely work with everyone of their members regularly throughout the competition, but we acknowledge this might be inappropriate for our questions regarding conflict experiences and learning experiences where some personal bias and subjectivity could occur.

The analysis and discussion of the study results possesses some limitations. The small body of literature on SynBio pedagogy made it difficult to evaluate the selected quotes in this report because the language to describe phenomena in SynBio education barely existed. Therefore, we needed to borrow and translate terminology from other disciplines into SynBio. Consequently, there are many instances where we introduce novel concepts in SynBio education that have not been described in literature pertaining to SynBio. This includes transdisciplinarity, the DiTO diagram, and the SNIS scoring rubric. These novel concepts are all grounded in existing literature in other disciplines and need to be studied further in the SynBio context, and tested/ validated by SynBio educators.

Lastly, we divided SynBio into the three dimensions WL, DL, and HP based on Canadian iGEM teams predominantly separating their work into sub-teams for each dimension. This division and terminology was intentionally not incorporated into the definition of SynBio that we provide at the beginning of this report because we acknowledged that there are some disciplines that fit between these dimensions, such as bioinformatics (WL-DL); and disciplines that don't fit neatly into these dimensions, such as prototyping and hardware development (considered DL in iGEM, but not involving any mathematical modelling). Consequently, we were not able to discuss the full breadth of material that could be taught in a SynBio curriculum, which deserves further attention.
